# Deubiquitination of phosphoribosyl-ubiquitin conjugates by PDE domain-containing *Legionella* effectors

**DOI:** 10.1101/745331

**Authors:** Min Wan, Alan Sulpizio, Anil Akturk, Wendy H.J. Beck, Michael Lanz, Vitor M. Faça, Marcus B. Smolka, Joseph P. Vogel, Yuxin Mao

## Abstract

Posttranslational protein modification by ubiquitin (Ub) is a central eukaryotic mechanism that regulates a plethora of physiological processes. Recent studies unveiled an unconventional type of ubiquitination mediated by the SidE family of *Legionella pneumophila* effectors, such as SdeA, that catalyzes the conjugation of Ub to a serine residue of target proteins via a phosphoribosyl linker (hence named PR-ubiquitination). Comparable to the deubiquitinases (DUBs) in the canonical ubiquitination pathway, here we show that two *Legionella* effectors, named DupA (<u>d</u>e<u>u</u>biquitinase for <u>P</u>R-ubiquitination) and DupB, reverse PR-ubiquitination by specific removal of phosphoribosyl-Ub (PR-Ub) from substrates. Both DupA and DupB are fully capable of rescuing the Golgi fragmentation phenotype caused by exogenous expression of SdeA in mammalian cells. We further show that deletion of these two genes results in significant accumulation of PR-ubiquitinated species in host cells infected with *Legionella*. In addition, we have identified a list of specific PR-ubiquitinated host targets and show that DupA and DupB play a role in modulating the association of PR-ubiquitinated host targets with *Legionella* containing vacuoles (LCV). Together, our data establish a complete PR-ubiquitination and deubiquitination cycle and demonstrate the intricate control that *Legionella* has over this unusual Ub-dependent posttranslational modification.

**Statement of significance:** Ubiquitination is a vital posttranslational modification in eukaryotes. A variety of microbial pathogens exploit this pathway during their infection. *Legionella pneumophila*, the causative bacterial pathogen of Legionnaires’ disease, has been show to hijack host ubiquitination pathway via a large number of effectors. Recent studies revealed a family of effectors catalyzing a novel type of Ub-dependent posttranslational modification, namely PR-ubiquitination. Here we report two new players, DupA and DupB, involved in this unconventional pathway. We found that DupA and DupB function as PR-Ub specific DUBs and play a role in regulating the PR-ubiquitination levels of host targets. Our results not only provide an expanding view of the PR-ubiquitination pathway, but may also facilitate the future identification of PR-ubiquitination pathways in eukaryotes.

## Introduction

Ubiquitin (Ub), a 76 amino acid protein, is attached to specific proteins as a potent posttranslational mechanism. Ubiquitination plays an essential role in a broad aspect of cellular processes, including protein homeostasis [1], cell signaling [2], and membrane trafficking [3, 4]. Following the conventional scheme of ubiquitination, Ub is covalently coupled to lysine residues on target proteins via the sequential activities of a collection of enzymes known as E1, E2, and E3s [5]. The C-terminal glycine residue of Ub is first activated and covalently linked to the catalytic cysteine residue of the Ub activating enzyme E1 through a thioester bond with the consumption of an ATP. The activated Ub moiety is then transferred to the active site cysteine of an E2 Ub-conjugating enzyme. The resulting thioester-linked E2∼Ub complex interacts with specific E3 Ub ligases, which promote the direct or indirect transfer of Ub to either the ε-amine of a lysine residue of targeted proteins or the N-terminal amine of another Ub molecule [6–8].

Given the vital role of ubiquitination in cell physiology, it is not surprising that a variety of microbial pathogens exploit this essential posttranslational modification pathway during the infection of their corresponding hosts [9]. For example, the intracellular pathogen *Legionella pneumophila* injects more than three hundred effector proteins into host cells via the Dot/Icm transporter [10, 11]. Among the hundreds of *L. pneumophila* effectors, more than 10 proteins are involved in ubiquitin manipulation [10]. These include proteins that contain the conserved eukaryotic F- or U-box domains found in some canonical E3 ubiquitin ligases [12–15]. Other E3 Ub ligases that have a unique structural fold but similar catalytic chemistry to the HECT-type ligases have also been characterized [16–18]. In addition to these Ub ligases, which utilize the canonical host Ub machinery for ubiquitination, recent studies of the *L. pneumophila* SidE family of effectors (SidEs), such as SdeA, uncovered a novel ubiquitination pathway that acts independently of E1 and E2 enzymes [19–21]. Instead, this unusual SdeA-catalyzed ubiquitination involves both mono-ADP-ribosyl transferase (mART) and phosphodiesterase (PDE) activities to PR-Ubiquitinate substrates. SdeA first uses its mART domain to catalyze the transfer of ADP-ribose from NAD^+^ to the residue R42 of Ub to generate mono-ADP-ribosyl Ub (ADPR-Ub). Subsequently, via the activity of the PDE domain, ADPR-Ub can be conjugated to serine residues of substrate proteins to generate a serine ubiquitinated product and release AMP (Fig. 1*A*). Alternatively, ADPR-Ub can be transferred to a water molecule to generate phosphoribosyl Ub (PR-Ub) in the absence of substrate proteins [22–26]. Our previous data further showed that the mART and the PDE domain function independently and the isolated PDE domain from SdeA is fully functional to PR-ubiquitinate substrates when ADPR-Ub is supplied [22].

**Fig. 1.**
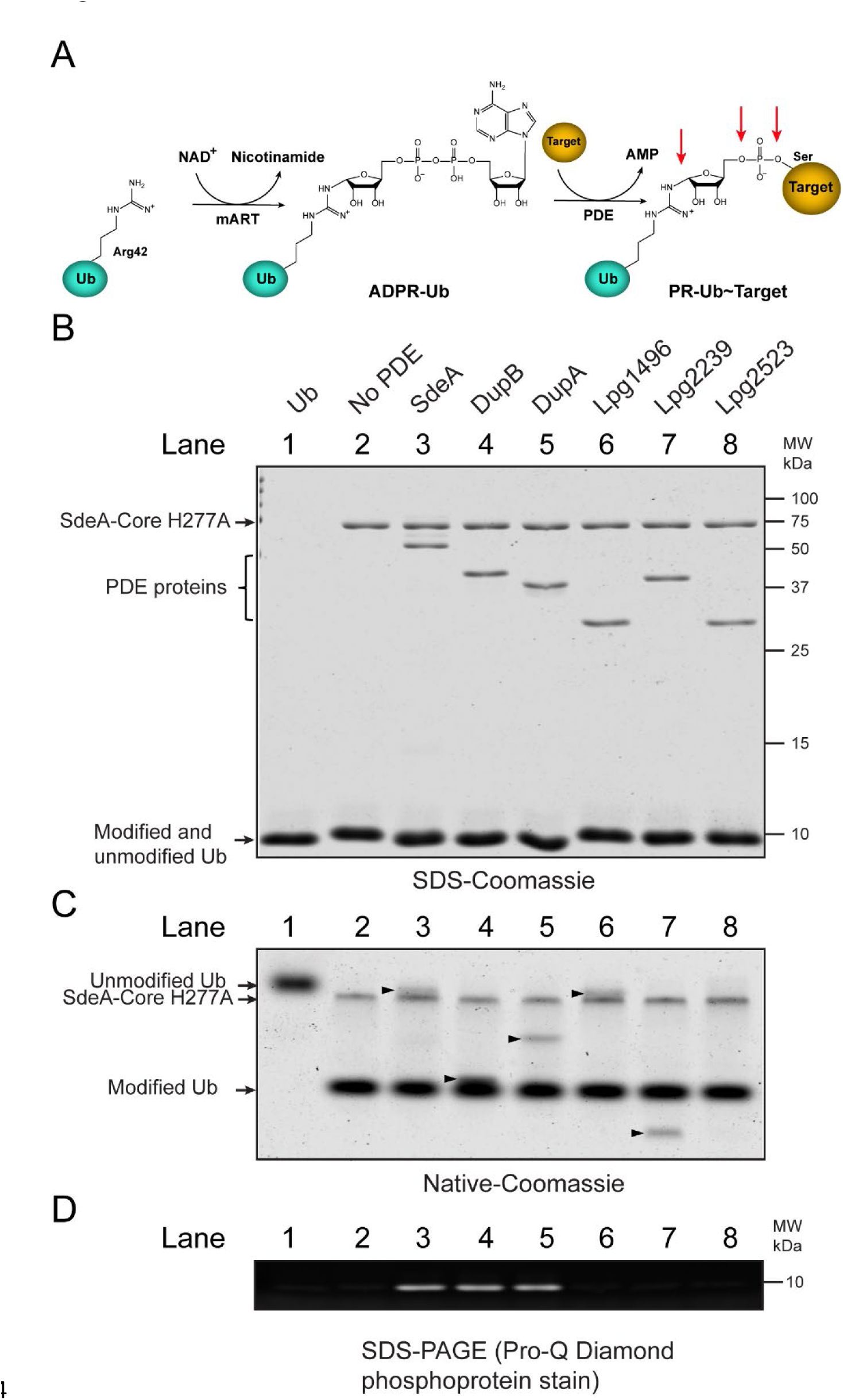
The PDE domains from DupA and DupB process ADPR-Ub to PR-Ub. (*A*) Schematic view of the two-step PR-ubiquitination reaction. The mART domain of SidEs catalyzes ADP-ribosylation of residue R42 of Ub and the PDE domain conjugates PR-Ub to a serine residue of substrates. Possible cleavage sites by potential PR-Ub specific DUBs are indicated by red arrows. (*B*) ADPR-Ub was first generated with SdeA-Core H277A and then mixed with the PDE domain from the indicated *Legionella* effectors. The final reaction mixtures were analyzed by SDS-PAGE followed by Coomassie Blue staining. The SdeA-Core, PDE, and Ub proteins are indicated on the left side of the gel. (*C*) Native PAGE analysis of the final reaction mixtures. The modification of Ub was exhibited as the mobility-shift of Ub species. The arrow heads mark the migration position of the PDE domain proteins on the native gel. (*D*) The processing of ADPR-Ub to PR-Ub was detected by SDS-PAGE followed by Pro-Q Diamond phosphoprotein staining. The signals in lane 3-5 indicate the production of PR-Ub.

The PDE domain of SdeA is conserved in nine *Legionella* effectors (*SI Appendix*, Fig. S1). Besides the three additional SdeA paralogs, SidE, SdeB, and SdeC, which have been shown to be PR-ubiquitination ligases, there are five other PDE domain-containing effectors that lack an adjacent mART domain. Here we report that the latter five PDE domain-containing proteins do not possess PR-Ub ligase activity, but surprisingly, Lpg2154 and Lpg2509 (previously named SdeD [27]) exhibit a robust activity to process ADPR-Ub to PR-Ub and efficiently cleave PR-Ub from PR-ubiquitinated substrates. We thus renamed Lpg2154 and Lpg2509 as DupA (<u>d</u>e<u>u</u>biquitinase for <u>P</u>R-ubiquitination) and DupB, respectively. We further show that DupA and DupB can rescue the dispersed Golgi phenotype caused by exogenous expression of SdeA in mammalian cells. Finally, we demonstrate that a strain lacking *dupA* and *dupB* markedly promotes the accumulation of PR-ubiquitinated species in host cells during infection. The presence of the two bona fide PR-Ub specific DUBs provides a potential regulatory mechanism for PR-ubiquitination in *Legionella* infection.

## Results

### Two PDE domain-containing *Legionella* effectors can process ADPR-Ub to generate PR-Ub

Our previous studies have shown that the isolated PDE domain of SdeA is sufficient to PR-ubiquitinate substrates when purified ADPR-Ub is supplied. Since the PDE domain is conserved in a total of nine effectors from the Philadelphia strain of *Legionella pneumophila* (*SI Appendix*, Fig. S1), we asked whether other PDE domains have similar PR-ubiquitination ligase activity as SdeA. To answer this question, we prepared recombinant proteins of the PDE domain from SdeA and five non-SidE family PDE-domain containing *Legionella* effectors. We then incubated the PDE domain proteins with purified HA-tagged ADPR-Ub and whole HEK293T cell lysates for 1 hour at 37 °C to allow the PR-ubiquitination reaction to occur. PR-ubiquitinated species were generated in the reaction with the PDE domain of SdeA but not any other PDEs (*SI Appendix*, Fig S2*A*). Consistent with these results, only the SdeA PDE domain could PR-ubiquitinate Rab33b, a previously reported PR-ubiquitinated substrate [19] (*SI Appendix*, Fig. S2*B*).

Since the PDE domain of SdeA is able to cleave ADPR-Ub to generate PR-Ub and AMP in the absence of substrates [20, 22], we asked whether other PDE domains are also able to process ADPR-Ub to PR-Ub. To address this question, we performed a two-step reaction to track the modification of Ub. In this assay, ADPR-Ub was first generated with a mutant SdeA fragment containing only an active mART domain (SdeA-Core H277A, amino acid 211-910). After the first step of the reaction, each PDE domain protein was then added to the reaction to determine their ability to process ADPR-Ub. All the proteins involved in the reactions were analyzed SDS-PAGE (Fig. 1*B*). The modification of Ub was indicated by the mobility shift of modified Ub in native-PAGE (Fig. 1*C*) and the generation of PR-Ub were analyzed by SDS-PAGE followed by Pro-Q Diamond phosphoprotein staining, which specifically stains terminal phosphoryl groups (Fig. 1*D*). Interestingly, in addition to the PDE domain of SdeA (Fig. 1*A-C*, lane 3), both the PDE domains from DupB and DupA were able to cleave ADPR-Ub to generate PR-Ub (Fig. 1*A-C*, lane 4 and 5). In contrast, the PDE domains of Lpg1496, Lpg2239, and Lpg2523 were not able to hydrolyze ADPR-Ub (Fig. 1*A-C*, lane 6 to 8). These results demonstrate that the PDE domains of DupA and DupB are able to process ADPR-Ub to PR-Ub, indicating DupA and DupB may modulate the PR-ubiquitination pathway during *Legionella* infection.

### DupA and DupB are PR-ubiquitination specific deubiquitinases

The cleavage of ADPR-Ub to PR-Ub by DupA and DupB led us to hypothesize that DupA and DupB may function as a deubiquitinase that specifically cleaves PR-Ub from PR-ubiquitinated substrates. To test this hypothesis, we incubated whole cell lysates from HEK293T cells with recombinant HA-Ub, SdeA-Core, and NAD^+^ to generate PR-ubiquitinated substrates. The PR-ubiquitinated products were then incubated with the indicated wild type and catalytically inactive mutant PDE domains. Strikingly, PR-ubiquitination signals were markedly reduced after incubation with the wild type PDE domains of DupA and DupB, but not their catalytically inactive mutants (Fig. 2*A*, lane 3-6). By contrast, the PDE domain of SdeA did not reduce the PR-ubiquitination signal (Fig. 2*A*, lane 2) even though it can process ADPR-Ub to PR-Ub (Fig. 1). Similar to SdeA, the other PDE-domain containing effectors, Lpg1496, Lpg2239 and Lpg2523 showed no effect on the levels of PR-ubiquitinated substrates (Fig. 2*A*, lane 7 to 9). Furthermore, the *L. pneumophila* effector SidJ, which was previously reported as a PR-Ub specific DUB [28], showed no detectable changes of the PR-ubiquitinated signals when the PR-ubiquitinated substrate mixtures were incubated with purified recombinant SidJ and Calmodulin (Fig. 2*A*, lane 10). To further validate this ability, we examined whether DupA and DupB could process a PR-ubiquitinated protein when they were expressed within cells. PR-ubiquitinated Rab33b was processed only by wild type DupA and DupB but not DupA or DupB catalytic mutants or other PDE domain-containing proteins, including once again SidJ (Fig 2*B* and *SI Appendix*, Fig. S3). Finally, to address whether DupA and DupB can specifically process only PR-Ub conjugates and not conventional Ub conjugates, we prepared whole cell lysates from cells expressing HA-Ub and then treated it with the canonical N-terminal DUB domain of SdeA or the PDE domain-containing proteins. The ubiquitination signals were drastically reduced in the presence of the SdeA DUB domain, however, no observable changes were detected in the presence of each PDE domain-containing proteins (*SI Appendix*, Fig. S4). Together, these data demonstrated that DupA and DupB are bona fide PR-Ub specific deubiquitinases.

**Fig. 2.**
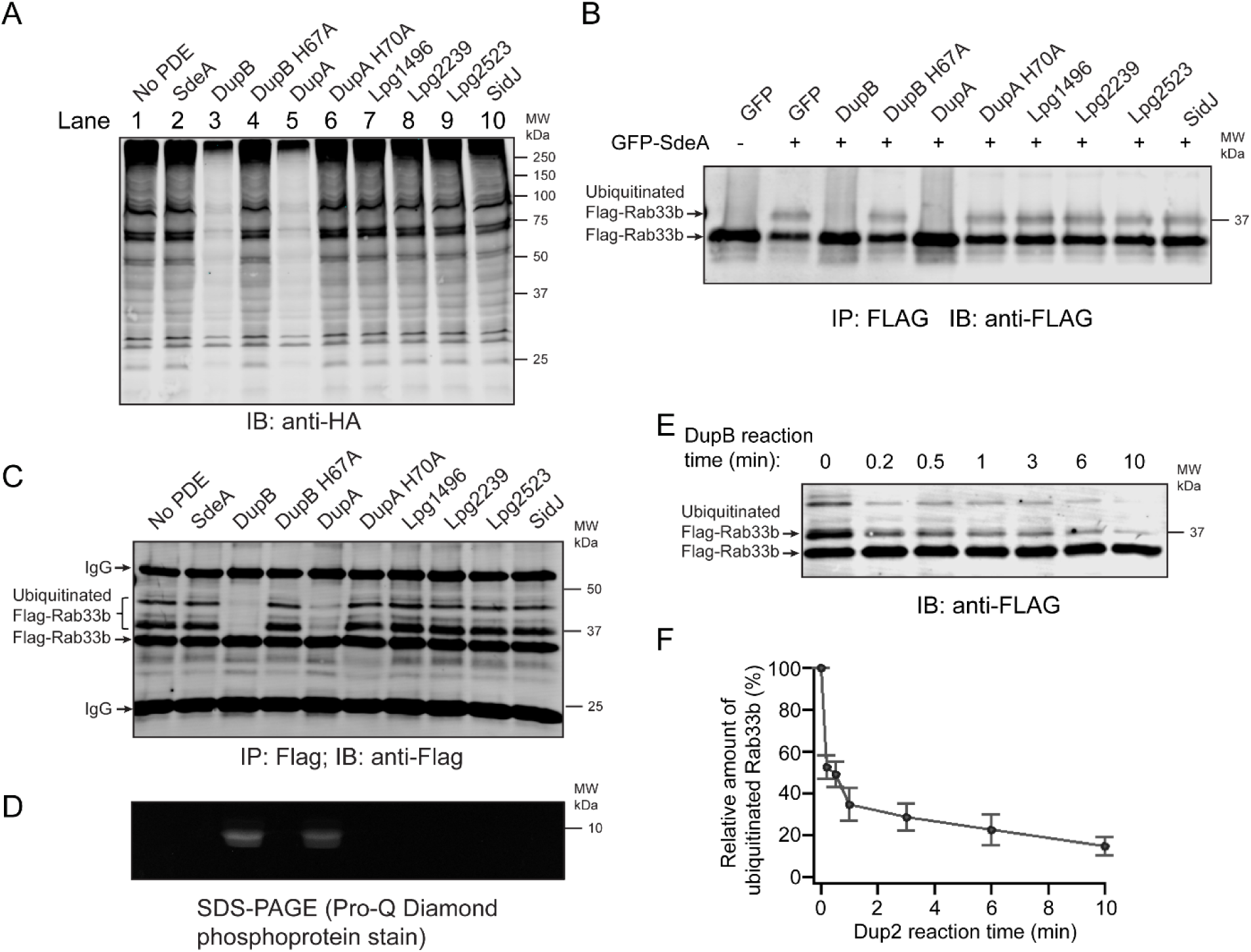
DupA and DupB function as PR-Ub specific deubiquitinases. (*A*) PR-ubiquitinated species were generated by incubating HEK293T cell lysates with HA-Ub, NAD^+^, and SdeA-Core. The PR-ubiquitinated products were then treated with recombinant proteins of the PDE domains from the indicated *Legionella* effectors or SidJ. The final reaction products were analyzed by Western blot against HA. (*B*) 4xFlag-Rab33b and EGFP-SdeA were co-expressed with EGFP-tagged full length *Legionella* effectors in HEK293T cells. 4xFlag-Rab33b was immunoprecipitated and analyzed by Western blot against anti-Flag. (*C*) Recombinant 4xFlag-Rab33b was PR-ubiquitinated by SdeA-Core and purified with anti-Flag beads. The bound proteins were then treated with the indicated recombinant PDE domain proteins or SidJ and the final products were analyzed by anti-Flag Western blot. (*D*) Pro-Q phosphoprotein staining of the final products from (*C*) to detect the generation of PR-Ub. (*E*) Reaction time course of the cleavage of PR-ubiquitinated Rab33b by DupB. PR-ubiquitinated Rab33b was generated using 6xHis-sumo-SdeA-Core. After removal of 6xHis-sumo-SdeA-Core with cobalt beads, the supernatant was treated with 1 μM DupB PDE domain for the indicated period of time at 37 °C. At each time point, equal amounts of reaction volumes were mixed with SDS loading dye to stop the reaction and analyzed by anti-Flag Western blot. (*F*) Quantification of the cleavage of PR-ubiquitinated Rab33b in (*E*). The intensity of the remaining PR-ubiquitinated Rab33b at each reaction time point was normalized to the value at time 0. The error bars represent the standard error of mean (SEM) of three independent experiments.

Since PR-Ub is linked to a serine residue of substrates via a ribose and phosphate group [19–21], we next investigated which bond is cleaved by DupA and DupB and what products are generated after the cleavage (see potential cleavage sites in Fig. 1*A*). To address this question, we generated PR-ubiquitinated Flag-Rab33b using SdeA-Core and immobilized it (unmodified or the PR-ubiquitinated form) onto beads conjugated with anti-Flag antibodies. The immunoprecipitated materials were then treated with PDE domain proteins or SidJ. The final products were analyzed by an anti-Flag Western blot and Pro-Q diamond phosphoprotein stain. In agreement with our previous results, the amount of PR-ubiquitinated Rab33b diminished after treatment with DupA and DupB, but not by their catalytically defective mutants or by any of the other PDE domain proteins (Fig. 2*C*). Importantly, phosphoprotein signals were detected in samples treated with wild type DupA or DupB and appeared at a band corresponding to the size of Ub, indicating the production of PR-Ub (Fig. 2*D*). Therefore, these results suggest that DupA and DupB cleave the phosphoester bond between the phosphate group and the hydroxyl group of the serine residue of the substrate to generate PR-Ub and unmodified substrates. The production of PR-Ub was further confirmed by mass spectrometry analysis (*SI Appendix*, Fig. S5). The cleavage of the phosphoester bond between the phosphate group and the hydroxyl group of the serine residue by DupA/DupB is also in agreement with our published crystal structure of the PDE domain of DupB in complex with ADPR-Ub, in which the β-phosphate group of ADPR-Ub is positioned and primed to be attached by the catalytic histidine (H67) to produce PR-Ub [22] (*SI Appendix*, Fig. S6). To further investigate the PR-Ub DUB activity of DupA/DupB, we performed semi-quantitative enzymatic assays by measuring the clearance rate of PR-ubiquitinated Rab33b in the presence of the DupB PDE domain (Fig 2*E*). The amount of PR-ubiquitinated Rab33b was reduced to less than 50% within 1 min of reaction when treated with DupB at 1 µM of concentration (Fig 2*E* and *F*). However, the levels of PR-ubiquitinated Rab33b remained unchanged when treated with the same amount of SdeA PDE or SidJ proteins under similar conditions (*SI Appendix*, Fig. S7).

### Golgi fragmentation caused by SdeA can be suppressed by DupA and DupB

It has been reported that exogenous expression of SdeA induces Golgi fragmentation in mammalian cells [29]. We were able to recapitulate the dispersed Golgi phenotype in HeLa cells when wild type full length SdeA was expressed. Furthermore, we found that both the intact mART and PDE activities were required to cause Golgi fragmentation since cells expressing the SdeA EE/AA (E860A/E862A) mutant, which is defective in mART activity, or the SdeA H277A mutant, which is defective in PDE activity, showed normal Golgi morphology (*SI Appendix*, Fig. S8). These observations suggest that the dispersed Golgi morphology is due to PR-ubiquitination of host proteins mediated by SdeA. The identification of DupA and DupB as PR-Ub specific DUBs led us to hypothesize that DupA and DupB might be able to suppress the dispersed Golgi phenotype caused by SdeA. Indeed, the Golgi morphology was restored when SdeA transfected cells were co-transfected with either mCherry-DupA or DupB, but not their PDE defective mutants or other PDE-domain containing effectors (Fig. 3). In agreement with previously reported results, the Golgi morphology was also rescued by co-transfection with the *Legionella* effector SidJ (Fig. 3). In summary, the dispersed Golgi caused by SdeA is likely due to PR-ubiquitination of host proteins and either DupA or DupB can fully restore the Golgi morphology by its PR-Ub specific DUB activity.

**Fig. 3.**
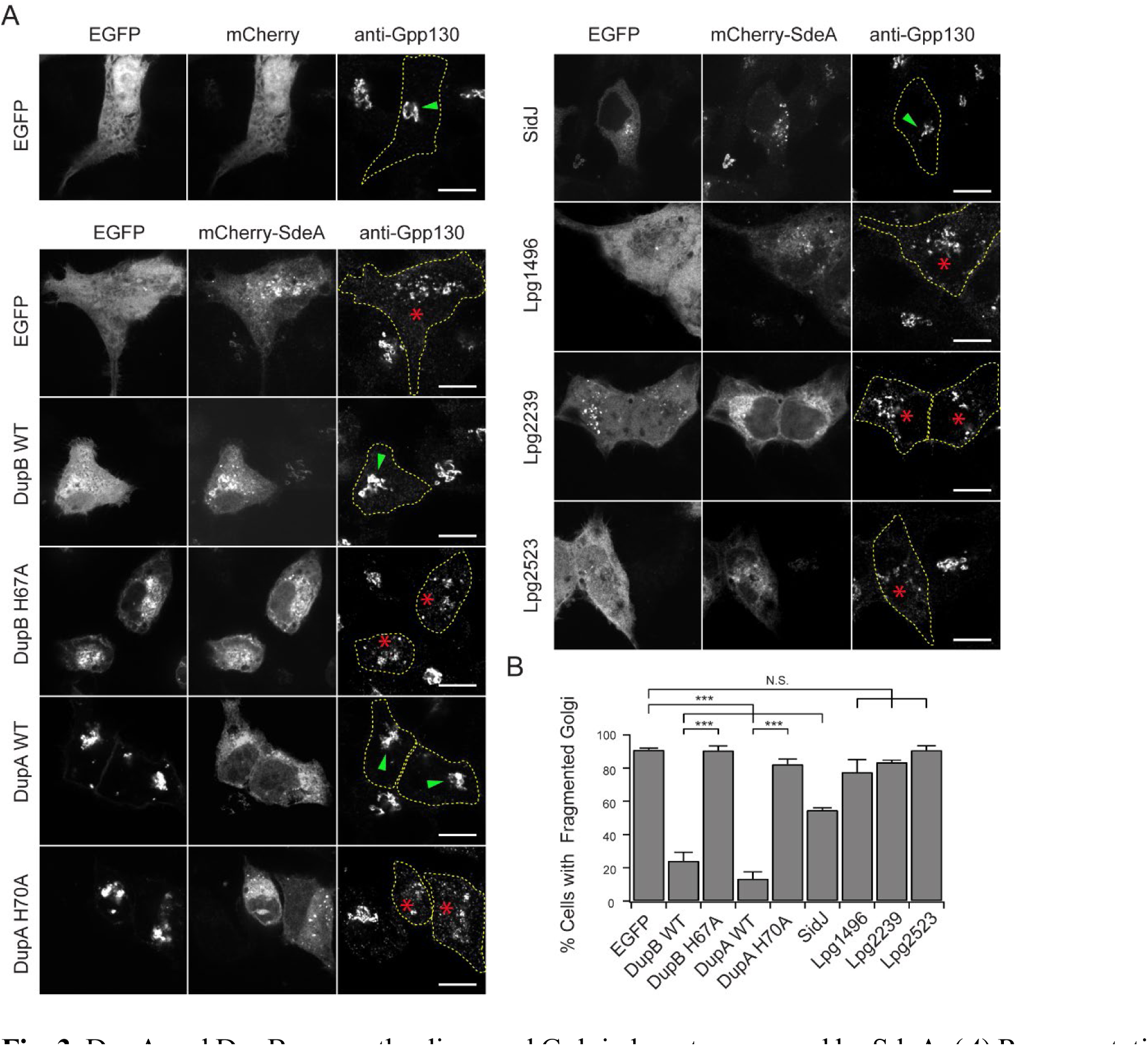
DupA and DupB rescue the dispersed Golgi phenotype caused by SdeA. (*A*) Representative images showing the Golgi morphology of HeLa cells co-expressing mCherry-SdeA and EGFP-tagged full length *Legionella* effectors. Cells were fixed and immunostained with rabbit anti-Gpp130. In transfected cells, cell outlines were drawn with dashed lines and Golgi structures with normal stacked or dispersed morphology were indicated with green arrow heads or red stars, respectively. Scale Bar: 10μm. (*B*) Percent of cells showing dispersed Golgi for the cells expressing the indicated proteins. Data are shown as means ± SEM of three independent experiments. At least 40 cells/condition were counted. ***P<0.001, N.S., not significant.

### DupA and DupB do not suppress SdeA toxicity in yeast

Previously, it was shown that expression of SdeA in yeast strongly inhibits its growth [19, 29–32]. Inhibition by SdeA requires its mART activity since the SdeA mART mutant (SdeA EE/AA) is no longer toxic. In contrast, SdeA’s DUB and PDE activities are not required for toxicity as a SdeA DUB mutant (SdeA C118A) and a SdeA PDE mutant (SdeA H277A) remained inhibitory (*SI Appendix*, Fig. S9*A-C*). Expression of the *Legionella* effector SidJ, which is a negative regulator of SdeA activity, was able to suppress SdeA toxicity when the proteins were co-expressed (*SI Appendix*, Fig. S9*D*) [28, 29]. As a result, we inquired whether DupA or DupB could similarly alleviate the toxicity of SdeA in yeast. In contrast to SidJ, DupA and DupB were co-expressed in yeast with SdeA (*SI Appendix*, Fig. S9*D* and *E*). Surprisingly, co-expression of DupA or DupB using either the *Pcyc* or *Pgal* promotors could not suppress the toxicity of SdeA (*SI Appendix*, Fig. S9D-E and Fig. S10). Therefore, even though DupA and DupB are bona fide PR-Ub specific DUBs, they are unable to block toxicity of SdeA in yeast suggesting a mechanistic difference between DupA/DupB and SidJ (discussed further below).

### DupA and DupB regulate PR-ubiquitination of host targets in *Legionella* infection

To test if DupA and DupB function as PR-Ub specific DUBs during intracellular infection by *L. pneumophila*, we created Δ*dupA*, Δ*dupB,* and `Δ`*dupA* `Δ`*dupB* double deletion mutants and we confirmed that *dupA* deletion does not interfere with the expression of *sidJ*, which is situated next to *dupA* in the *L. pneumophila* genome (*SI Appendix*, Fig. S11). We theorized the absence of DupA and/or DupB would result in the over-accumulation of PR-ubiquitinated proteins during an infection. As predicted, infection of `Δ`*dupA* or `Δ`*dupB* single knockout strains resulted in a modest increase in PR-ubiquitinated Rab33b in HEK293T cells whereas substantially more PR-ubiquitinated Rab33b accumulated using the `Δ`*dupA* `Δ`*dupB* double knockout strain (Fig. 4*A*, lane 1-5). This effect was due to the absence of DupA and DupB as the double mutant could be complemented by expression of wild type DupA or DupB, but not their catalytic mutants (Fig. 4*A*, lane 6 to 9). Furthermore, the levels of PR-ubiquitinated Rab33b remained at a low level up to 6 hours post infection in the WT strain while the PR-ubiquitinated Rab33b levels increased dramatically in the `Δ`*dupA* `Δ`*dupB* mutant over the same time frame of infection (Fig. 4*B*). The elevated levels of PR-ubiquitinated substrates were not limited to the exogenously overexpressed substrate Rab33b. The enhancement of PR-ubiquitination of endogenous host proteins was also evident in cells infected with the `Δ`*dupA* `Δ`*dupB* strain (Fig 4*C*). These results convincingly demonstrate that DupA and DupB play a role in regulating the PR-ubiquitination of host targets during *Legionella* infection. We next asked whether the deletion of these two PR-Ub specific DUBs has an effect on *Legionella* intracellular growth. Unlike the Δ*sidJ* mutant, the Δ*dupA ΔdupB* mutant does not have a significant growth defect on its own within *Acanthamoebae castellanii*. However, similar to what was previously seen with a Δ*sidJ* mutant [29], the growth of the Δ*dupA ΔdupB* mutant was inhibited by over-expression of SdeA (*SI Appendix*, Fig. S12). These data suggest a mechanistic distinction between DupA/DupB and SidJ and further support a role of DupA and DupB in regulating the PR-ubiquitination pathway.

**Fig. 4.**
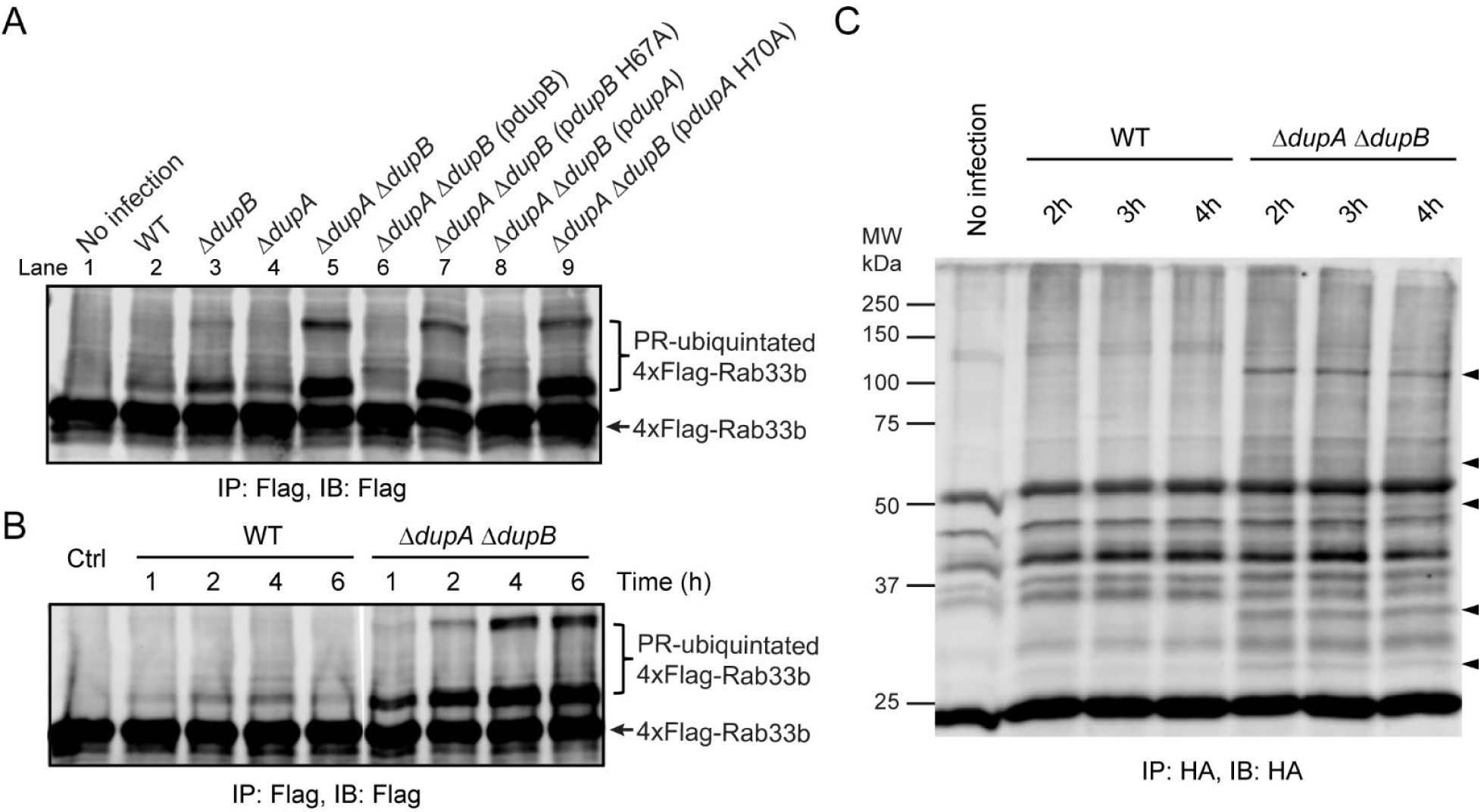
DupA and DupB regulate PR-ubiquitination of host targets during *Legionella* infection. (*A*) HEK293T cells expressing 4xFlag-Rab33b were infected with relevant *L. pneumophila* strains for 2 hours. Rab33b proteins were enriched by anti-Flag immunoprecipitation and analyzed by anti-Flag Western blot. (*B*) HEK293T cells expressing 4xFlag-Rab33b were challenged with either WT or the `Δ`*dupA* `Δ`*dupB* strain for the indicated periods of time and Rab33b proteins were immunoprecipitated and analyzed by anti-Flag Western blot. (*C*) HEK293T cells expressing HA-Ub G76A were infected with WT or `Δ`*dupA* `Δ`*dupB Legionella* strains. PR-ubiquitinated host proteins were enriched by anti-HA immunoprecipitation and analyzed by anti-HA Western blot. Specific bands detected under the infection by Δ*dupA* Δ*dupB* strain were marked with arrow heads. For all the infection experiments, cells were co-transfected with a plasmid expressing FcγRII and the *Legionella* bacteria were opsonized by polyclonal *Legionella* antibodies prior to infection.

### Identification of PR-ubiquitinated host targets in *Legionella* infection

The increased levels of PR-ubiquitinated proteins were apparent during an infection with the `Δ`*dupA* `Δ`*dupB* mutant provided us with the opportunity to identify PR-ubiquitinated host targets via a SILAC (Stable Isotope Labeling by Amino acids in Cell culture) approach (Fig. 5*A*). In this approach we employed a Ub mutant, HA-Ub G76A, as it has a decreased propensity to be used in the conventional Ub pathway, but can be used in the PR-ubiquitination pathway due to its intact residue R42. We transformed HEK293T cells with HA-Ub G76A and then challenged the cells grown in medium containing light isotope labeled Arg/Lys with the `Δ`4*sidE* strain, which lacks all four members of the SidE family, and cells grown in medium containing heavy isotope labeled Arg/Lys with the `Δ`*dupA* `Δ`*dupB Legionella* strain for 2 hours. Cell lysates were prepared, treated, and analyzed by LC-MS/MS (Fig. 5*A*). Applying these strategies, we identified a list of potential host PR-ubiquitination targets by mass spectrometry analysis (Fig 5*B* and *SI Appendix*, Table S1). In agreement with a previous publication [21], we identified the endoplasmic reticulum (ER) protein Rtn4 as a PR-ubiquitinated protein. Other substrates included GORASP2 (Golgi reassembly-stacking protein 2, also named GRASP55) and TOR1AIP2 (Torsin-1A-interacting protein 2, also named LULL1). GRASP55 is a Golgi and ER membrane localized protein and plays a role in membrane stacking of Golgi cisternae [33, 34] or in regulating an unconventional ER-stress induced secretory pathway [35]. The other protein, LULL1 is an ER membrane protein regulating the activity and distribution of a AAA family ATPase, Torsin 1A, between ER and the nuclear envelope [36, 37]. We validated this approach using both GRASP55 and LULL1. PR-ubiquitinated LULL1 could be detected in cells infected with the wild type *L. pneumophila* strain Lp02 and a more robust signal were observed in cells infected with the `Δ`*dupA* `Δ`*dupB* mutant. However, no PR-ubiquitinated LULL1 was detected in cells in the absence of infection or cells infected with the `Δ`*4sidE Legionella* strain, which lacks PR-ubiquitination activity (Fig 5*C*). Similarly, PR-ubiquitinated GRASP55 was also confirmed in cells infected with wild type *L. pneumophila* strain Lp02 and the `Δ`*dupA* `Δ`*dupB* mutant (*SI Appendix*, Fig. S13*A*). Thus, this approach has revealed with high confidence a set of host targets that are PR-ubiquitinated during *Legionella* infection.

**Fig. 5.**
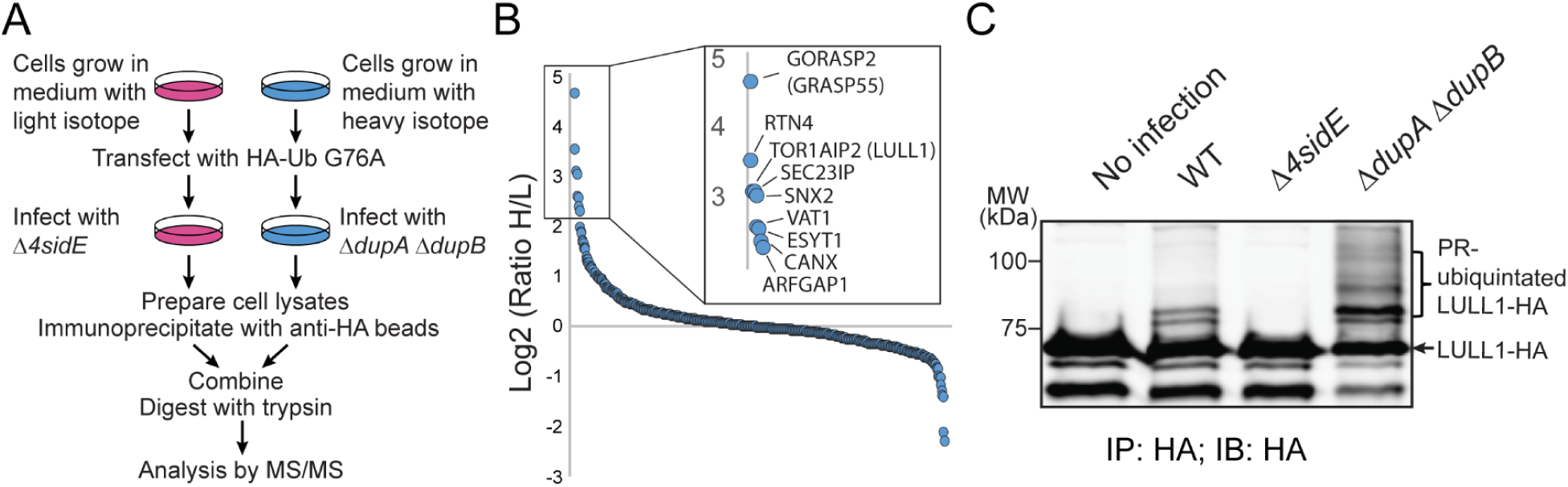
Proteomic identification of host PR-ubiquitination targets. (*A*) Experimental flow chart of SILAC sample preparation. (*B*) MS/MS analysis of SILAC samples prepared from cells expressing HA-Ub G76A and infected with either `Δ`*4sidE* (Light) or `Δ`*dupA* `Δ`*dupB* (Heavy) strains. The geometric mean of Heavy/Light peptide ratios for all detected proteins are plotted from left to right in descending order. For a protein to be considered in the plot, it required SILAC quantification from at least 6 independent peptide identifications. (*C*) Confirmation of PR-ubiquitinated host targets. HEK293T cells were transfected with LULL1-HA and then infected with relevant *Legionella* strains for 2 hours. LULL1 was enriched by anti-HA immunoprecipitation and analyzed by anti-HA Western blot.

### DupA and DupB modulate the recruitment of PR-ubiquitinated host proteins to *Legionella* containing vacuoles

Since PR-ubiquitinated Rtn4 was recruited to the *Legionella* containing vacuole (LCV) and the Rtn4-LCV association depended on the PR-ubiquitination activity of SidEs [21, 38], we hypothesized that other PR-ubiquitinated host substrates may also be recruited to the LCV. We tested this hypothesis by examining the location of LULL1-HA and GRASP55-HA in HEK293T cells infected with several different *Legionella* strains. In cells infected with wild type *L. pneumophila*, 49% of the LCVs were positive for LULL1 two hours post infection (Fig. 6*A* and *B*). In contrast, LULL1 recruitment did not occur with a Dot/Icm deficient strain or the `Δ`*4sidE* strain. However, the association of LULL1 with the LCVs was nearly doubled in cells challenged with the `Δ`*dupA* `Δ`*dupB Legionella* strain. Importantly, enhanced LULL1 association did not occur when the `Δ`*dupA* `Δ`*dupB* mutant was complemented with wild type DupB but remained at high levels in cells expressing the DupB catalytically inactive H67A mutant (Fig. 6*A* and *B*). Interestingly, quantitative analysis of Lull1 signals at the LCV revealed that the fluorescence intensity of LCV-associated LULL1 was nearly doubled in cells infected with `Δ`*dupA* `Δ`*dupB* or `Δ`*dupA* `Δ`*dupB* + pDupB H67A strain compared to that in cells infected with WT or `Δ`*dupA* `Δ`*dupB +* pDupB strain (Fig 6*C*). This observation is apparently in agreement with increased levels of PR-ubiquitination substrates in cells infected with `Δ`*dupA* `Δ`*dupB* strain (Fig. 4). Furthermore, as expected, we also observed a similar pattern of GRASP55 recruitment to the LCV in cells upon *Legionella* infection (*SI Appendix*, Fig. S13*B*-*D*). Therefore, PR-ubiquitination of more than one protein results in LCV-association during an infection and the PR-Ub DUB activity of DupA and DupB plays a role in modulating the recruitment of PR-ubiquitinated host targets to the bacterial phagosome.

**Fig. 6.**
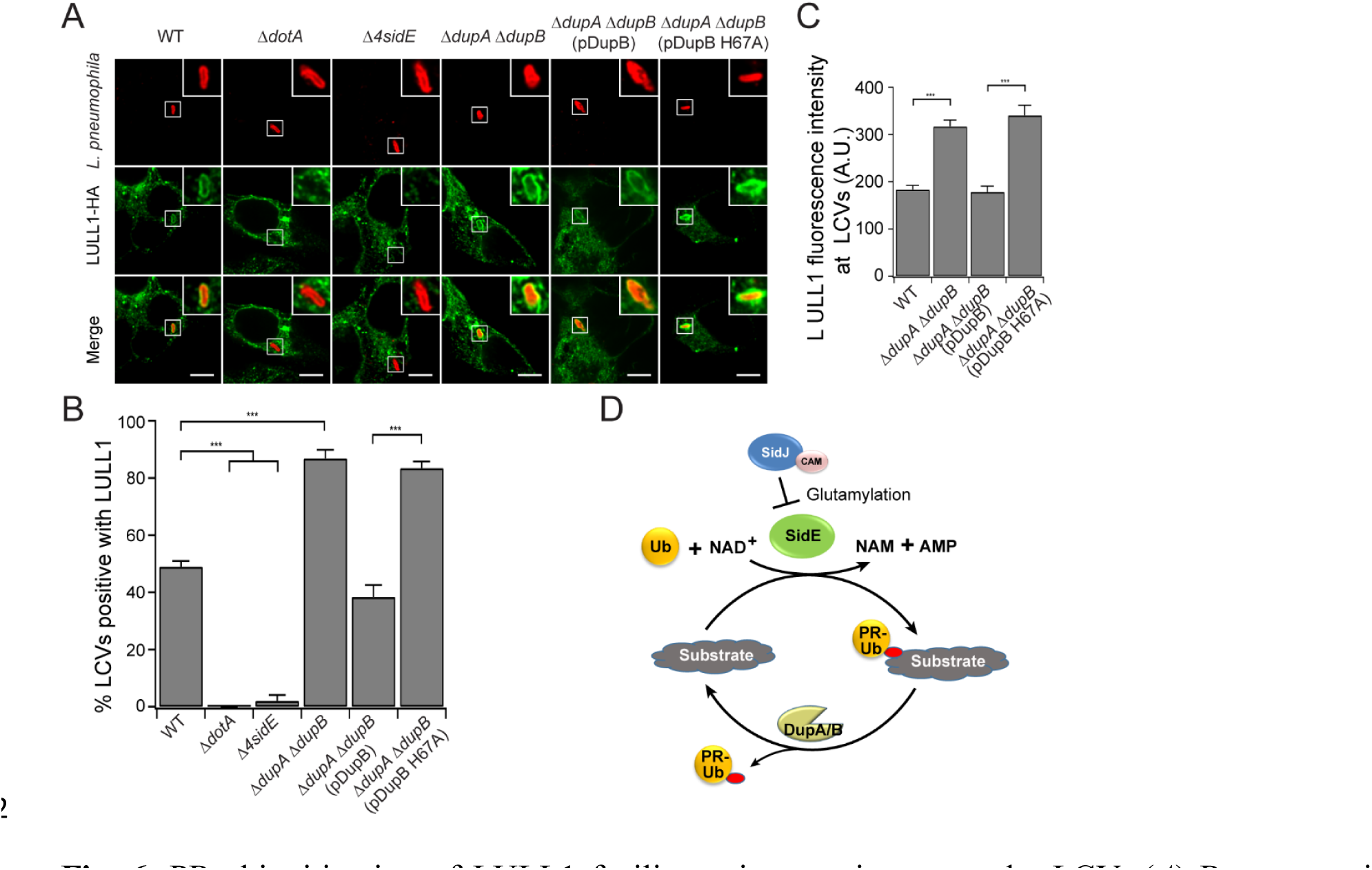
PR-ubiquitination of LULL1 facilitates its recruitment to the LCV. (*A*) Representative confocal images show the recruitment of LULL1 (green) in HEK293T cells challenged with specified *Legionella* strains (red) for 2 hours. Scale Bar: 5 μm. (*B*) Quantitative analysis of the percentage of LULL1-positive LCVs in cells infected with the indicated *Legionella* strains. Data are shown as means ± SEM of three independent experiments. More than 80 LCVs were counted for each condition. ***P<0.001. (*C*) Quantitative analysis of the LULL1 fluorescence intensity associated with the LCV. Data are shown as means ± SEM of three independent experiments. The value was averaged from more than 35 LULL1-positve LCVs for each condition. ***P<0.001. (*D*) Schematic model of the PR-ubiquitination and deubiquitination cycle. The SidE family PR-Ub ligases catalyze PR-ubiquitination while DupA and DupB catalyze the removal of PR-Ub from substrates. The effector SidJ inhibits the mART activity of SidE ligases by polyglutamylation modification of SidE family ligases in a Calmodulin (CAM)-dependent manner.

## Discussion

In our search for additional PR-ubiquitination ligases among *Legionella* effectors, we discovered two PDE domain-containing effectors, DupA (Lpg2154) and DupB (Lpg2509) function as PR-Ub specific deubiquitinates (DUBs). We showed that DupA and DupB can specifically process ADPR-Ub to PR-Ub and can deubiquitinate PR-ubiquitinated substrates to release PR-Ub and unmodified substrates. We also demonstrated that DupA and DupB are capable of rescuing the Golgi fragmentation phenotype caused by exogenous expression of SdeA in mammalian cells and can modulate the levels of PR-ubiquitination of endogenous host targets during infection. Our findings establish a complete PR-ubiquitination and deubiquitination cycle, in which the SidE family effectors catalyze the ligation of PR-Ub to substrates while DupA and DupB deubiquitinate PR-Ub from modified substrates.

Although we convincingly demonstrated that DupA and DupB function as PR-Ub specific DUBs, a recent study reported that the effector SidJ catalyzes the same PR-Ub deubiquitination reaction [28]. However, in contrast to this earlier report, we have been unable to detect this activity for SidJ. Consistent with our observed differences between SidJ and DupA/DupB, expression of SidJ, but not DupA or DupB, is able to suppress SidE-mediated yeast toxicity [29, 30, 39]. This is notable as this form of yeast toxicity can be recapitulated with the SdeA H277A mutant, which generates ADPR-Ub through its intact mART domain but lacks the PR-Ub ligase activity. As a result, yeast toxicity is not caused by PR-ubiquitination of host proteins but instead is likely due to the accumulation of ADPR-Ub or PR-Ub or depletion of Ub in yeast. Therefore, the most parsimonious explanation is that SidJ has a different activity other than as a PR-Ub DUB. Indeed, three recent publications reported that SidJ is polyglutamylase that attaches glutamate residues to SdeA and thus inhibits the mART activity of SdeA [40–42].

Our new results on DupA/DupB, as well as recently published data on SidJ, has revealed additional regulatory layers for how *Legionella* controls the PR-ubiquitination cycle during an infection (Fig. 6*C*). This pathway contains many factors including multiple PR-Ub ligases, PR-Ub specific DUBs, and PR-UB ligase inhibitors. The various PR-UB ligases (SidE, SdeA, SdeB, SdeC) presumably have different targets with the host cell and we speculate that the initial PR-ubiquitination triggered by these factors is important for the bacterium to establish a proper niche for intracellular growth. However, unregulated modification of the PR-UB ligase targets is toxic to the cell and therefore must be restricted. This could happen in multiple-independent steps. First, DupA and DupB function to remove a portion of the PR-ubiquitin tags from target proteins simultaneously with SidE ligases as the enhancement of PR-ubiquitinated substrates was evident as early as 1 hour post infection by the `Δ`*dupA* Δ*dupB* stain (Fig. 4*B*), thereby possibly limiting excessive PR-ubiquitination by the SidE ligases at the early infection stage. At a later point in the infection, SidJ directly suppresses the activity of SidE ligases after they are no longer needed consistent with the previously published observation that SidJ regulates the disappearance of SidE proteins from the LCV [29]. Finally, DupA and DupB may also act at later time points of infection to reverse the PR-ubiquitination of target proteins, thereby acting in concert with SidJ to down regulate the process.

Since all of the PDE domains from *L. pneumophila* effectors have a similar structural fold and nearly identical catalytic residues, it is puzzling that the PDE domains of the SidE proteins function as a PR-Ub ligase, whereas the PDEs domains of DupA and DupB deconjugate PR-ubiquitinated substrates. One possible explanation is that NMR chemical shift titration experiments revealed that the SdeA PDE domain showed no detectable interaction with Ub in solution, while the DupB PDE domain exhibited a direct and specific interaction with Ub [22]. As a result, it is possible that the differential Ub-binding affinity of the PDE domains determines whether they act as a ligase or DUB. In the case of SdeA, its PDE domain might bind ADPR-Ub, primarily due to the interaction with the ADPR moiety. However, its low Ub-binding affinity prohibits its binding with PR-ubiquitinated substrates, thus the PDE domain of SdeA can only function as a ligase but not a PR-Ub DUB. On the other hand, the PDE domains of DupA/DupB have a high affinity for Ub, which allows high affinity binding with both ADPR-Ub and PR-ubiquitinated substrates, thereby mediating their PR-Ub DUB activity. Nevertheless, future structural and biochemical studies will be required to resolve the molecular mechanism of different PDE domains.

Finally, by using the `Δ`*dupA* `Δ`*dupB Legionella* mutant. we were able to identify additional host PR-ubiquitinated substrates with a SILAC approach and demonstrated that modified LULL1 or GRASP55 was recruited to the LCV. Interestingly, many of the top hits are proteins localized on the endoplasmic reticulum, the Golgi apparatus, or intermediate transport vesicles between these membrane-bound organelles. This is consistent with the previous observation that PR-ubiquitination of Rtn4 is required for its recruitment to the LCV [21]. Therefore, PR-ubiquitination may play a key role in hijacking host endomembrane system during the maturation of bacterial containing vacuoles. Another interesting finding of our SILAC experiment is that quite a few proteins involved in canonical ubiquitination pathways are also among the top hits of the list. These proteins include the Ub activating enzyme E1, UBA1; Ub-conjugating enzyme E2s, UBE2L3 and UBE2; several Ub E3 ligases; and Ub adaptor proteins, Bag6 and p62/SQSTM1. Although these targets have yet to be confirmed under infection conditions, we speculate that PR-ubiquitination of these proteins may inhibits host endogenous ubiquitination pathways and thus may interfere with some Ub-dependent cellular process, such as autophagy, for the establishment of bacterial containing vacuoles. In addition, it is plausible that inhibition of host endogenous ubiquitination pathways diverts additional free Ub for the usage by the SidE family PR-Ub ligases. The exact biological consequence of PR-ubiquitination remains largely unknown, nevertheless, the list of potential PR-ubiquitinated substrates generated by using the `Δ`*dupA* `Δ`*dupB Legionella* mutant will certainly serve as a stepping stone for future investigations of this unusual type of Ub-dependent posttranslational modification.

To date, the PR-ubiquitination pathway is only found in *Legionella* species. It remains a fascinating question whether other organisms, particularly eukaryotes, encode a similar unique Ub-dependent posttranslational modification mechanism. The identification of two novel PR-Ub specific DUBs not only completes the components involved in the PR-ubiquitination pathway but also provides key tools to probe for PR-ubiquitinated proteins and may facilitate the identification of PR-ubiquitination pathways in other organisms.

## Materials and Methods

The materials and methods are described at length in *SI Appendix, Materials and Methods*. This includes cloning, site-directed mutagenesis, expression and purification of recombinant proteins in vitro PR-ubiquitination assays, *Legionella* infection analyses, SILAC experiments, and immunofluorescence imaging.

## ACKNOWLEDGEMENT

We thank Dr. Zhao-Qing Luo (Purdue University) for *Legionella* strains and antibodies. Dr. Anthony Bretscher (Cornell University) for yeast strains, plasmids, and confocal microscope support. Dr. William Brown (Cornell University) for antibodies. This work was supported by National Institutes of Health (NIH) Grants 5R01GM116964 (Y.M.), R01GM097272 (M.B.S.).

## AUTHOR CONTRIBUTIONS

M.W., A.S., A.A, and Y.M. conceived the project. M.W., A.S., A.A, W.H.J.B., M.L. V.M.F. performed the experiments. M.W., A.S., M.B.S., J.P.V, and Y.M. analyzed the data. M.W., A.S., and Y.M. wrote the paper.

## Supplementary Materials

### Supplemental Materials and Methods

#### Cloning and Mutagenesis

For protein expression in *E. coli*, SdeA core, Ub, HA-Ub, 4xFlag-Rab33b were generated as described previously [1]. DNA fragments corresponding to the PDE domains of DupA (residue 1-333), DupB (residue 1-356), Lpg1496 (298-598), Lpg2239 (135-524), and Lpg2523 (494-751) were PCR amplified from genomic DNA of *Legionella pneumophila* Philadelphia strain and digested with BamH1 and XhoI. SidJ (89-853) was amplified similar to other effectors, but digested with BamH1 and SalI. Calmodulin was amplified from the pEGFP-C1 Calmodulin2 gene obtained from Dr. Zhao-qing Luo (Purdue University) and digested with BamH1 and Xho1. These DNA fragments were inserted into pET28a-6xHis-Sumo vector digested with the same restriction enzymes [2]. DupA and DupB single mutations were generated by site-directed mutagenesis.

For expression in mammalian cells, EGFP-SdeA [1], 4xFlag-Rab33b [3], and HA-Ub [4] were generated as described previously. For expression of PDE domain-containing proteins in mammalian cells, full length DupA, DupB, Lpg1496, Lpg2239, and Lpg2523 were amplified from genomic DNA of *Legionella pneumophila* Philadelphia strain. DNA fragments corresponding to full length DupA, DupB, Lpg1496, and Lpg2523 were digested with BamHI and XhoI while Lpg2239 was digested with BglII and XhoI. The digested DNA fragments were inserted into mCherry-C1 or EGFP-C1 vectors digested with BglII and SalI. LULL1-HA [5] was a generous gift from Dr. Christian Schlieker (Yale School of Medicine). GRASP55-HA, DNA fragment of GRASP55-HA was generated by PCR from GRASP55-GFP (from Dr. Yanzhuang Wang, University of Michigan), digested with BamHI and SalI and inserted into pcDNA3 vector digested with BamHI and XhoI.

For yeast expression, DNA fragments corresponding to SdeA point mutants were digested with BamH1 and Xho1 and cloned into the BM272 pGal vector digested with BamH1 and SalI. Full length DupA, DupB and SidJ were cloned into the pRS415 pGal HA or pRS415 pCyc HA plasmids with standard restriction cloning using the BamH1 and Xho1 or SalI restriction sites.

For *Legionella* expression, DNA fragments of full length DupA, DupB, and their PDE mutants were PCR amplified and digested with BamH1 and Xho1. The digested DNA fragments were inserted into pZL507 plasmid (gift from Dr. Zhao-Qing Luo Purdue University). For *Legionella* in-frame deletions, DNA fragments of 1.2 Kb upstream and 1.2 Kb downstream of the *dupA* gene were PCR amplified and cloned into the pSR47s vector (obtained from Dr. Zhao-Qing Luo).

### Protein expression and purification

Plasmids encoding the PDE domains were transformed into *E. coli* Rosetta cells. Single colonies were cultured in Luria-Bertani medium supplemented with 50 μg/ml kanamycin or 100 μg/ml ampicillin to mid-log phase. Protein expression was induced with 0.2 mM isopropy1β-D-1-thiogalactopyranoside (IPTG) overnight at 18°C. Harvested cells were resuspended in a buffer containing 150 mM NaCl, 20 mM Tris-HCl pH 8.0 and lysed by sonication. The lysate was clarified by centrifugation at 31,000*g* for 30 min at 4 °C, then supernatant was incubated with cobalt resin (Gold-Bio) for 2 hrs at 4 °C. The bead slurry was extensively washed with lysis buffer on a gravity column. The SUMO-specific protease Ulp1 was then added to the resin slurry to release the protein from the 6xHis-Sumo tag. The proteins of interest were then collected for further purification with size exclusion chromatography. The preparation of Ub, HA-Ub, ADPR-Ub, and HA-ADPR-Ub was performed according to the protocol previously described [1].

### Cell culture and transfection

HeLa and HEK293T cells were cultured in Dulbecco’s modified Eagle’s medium (CellGro) supplemented with 10% fetal bovine serum (Gibco) and 1% penicillin-streptomycin (Invitrogen) at 37 °C and 5% CO2. Transfection was performed using polyethyleneimine (PEI) reagent.

### PR-ubiquitination and deubiquitination assays

PR-ubiquitination reactions were performed by mixing 1 μM of SdeA-core PDE mutant with 25 μM ubiquitin in reaction buffer (50 mM NaCl and 50 mM Tris pH 7.5) for 1 hr at 37 °C and then indicated PDE domain proteins (to a final 1 μM of concentration) were added to the reaction mixture for another 1 hr at 37 °C. The final reaction products were assessed using both 12% SDS–PAGE and 8% native PAGE. SDS– PAGE gels were stained with Coomassie for the visualization of all proteins in the reaction and with Pro-Q Diamond phosphoprotein stain (Invitrogen) for the visualization of PR-Ub production. Native gels were stained with Coomassie for mobility shift of modified Ub. ADPR-Ub and PR-Ub migrate to the same position on a native gel (labeled as modified Ub), however, only PR-Ub is visible by Pro-Q phosphoprotein stain due to its terminal phosphoryl group [6].

For PR-Ub DUB assay, 4 μM of recombinant 4xFlag-Rab33b was first incubated with 1 μM SdeA-Core, 25 μM Ub, and 1 mM NAD^+^ for 1hr at 37 °C. 4xFlag-Rab33b was purified with anti-Flag beads. The bound proteins were then treated with recombinant proteins of PDE domain from the indicated *Legionella* effector or SidJ (Since SidJ requires Calmodulin to function, we also included 5 μM Calmodulin, 5 mM MgCl2, 5 mM Glutamate, 1 mM ATP in the reaction) for 1 hr at 37 °C. 4xFlag-Rab33b was analyzed by Western blot using anti-Flag antibodies after treatment with indicated PDE domain proteins or SidJ reaction mixture. The reaction products were analyzed using SDS–PAGE followed by Western blot with an anti-Flag antibody (Sigma-Aldrich) at a 1:2,500 dilution.

To assay the intracellular PR-ubiquitination of Rab33b, plasmids expressing 4xFlag-Rab33b, EGFP-SdeA, and full-length protein of indicated EGFP-tagged *Legionella* effector in HEK293T cells. Whole cell lysates were subjected to immunoprecipitation with anti-Flag beads and the products were analyzed using anti-Flag Western blot. The expression of EGFP–SdeA constructs was analyzed by Western blot using a polyclonal anti-GFP antibody.

### Antibodies, immunostaining, and fluorescent microscopy

Rabbit polyclonal anti-GFP antibodies were obtained from Dr. Anthony Bretscher (Cornell University) [7]. Anti-SdeA was obtained from obtained from Dr. Zhao-qing Luo (Purdue University). Anti-*L. pneumophila*, anti SidJ and anti-ICDH (isocitrate dehydrogenase) antibodies were described previously [8]. Anti-Gpp130 (BioLegend). anti-HA (Sigma-Aldrich), anti-Flag (Proteintech) antibodies were purchased commercially. Anti-*L. pneumophila* antibodies were described previously [9]. For Gpp130 staining, transfected HeLa cells were fixed using 4% PFA and staining with rabbit-anti-Gpp130 primary antibody (1:500) followed by Alexa Fluor donkey anti-rabbit 647 secondary antibodies. For *Legionella* bacterium staining, infected cells expressing LULL1-HA were fixed using 4% PFA. Extracellular bacteria labeled with rabbit polyclonal anti-*L. pneumophila* antibodies were stained with Alexa Fluor donkey anti-rabbit 647 secondary antibodies. The cells were then permeabilized with 0.1% triton and stained using mouse anti-HA antibody (1:1000) and then with donkey anti-rabbit 568 and donkey anti-mouse 488 secondary antibodies. To measure LULL1-HA or GRASP55-HA signals at the LCV, a 10-pixel (∼0.54 μm) wide free hand circular line was drawn along the surface of each LCV using the ImageJ software (NIH). The LULL1 or GRASP55 fluorescence intensity was calculated as the mean fluorescence along the line for each LCV.

Images were acquired on a spinning disk confocal microscope (Intelligent Imaging Innovations, Denver, CO) with a spinning disk confocal unit (Yokogawa CSU-X1), an inverted microscope (Leica DMI6000B), a fiber-optic laser light source, a 100× 1.47NA objective lens, and a Hamamatsu ORCA Flash 4.0 v2+ sCMOS camera. Images were acquired using SlideBook 6.0 software and analyzed using ImageJ software.

### Immunoprecipitation and Western Blot

To analyze PDE-domain containing proteins and PR-ubiquitination substrate Rab33b, LULL1, or GRASP55, the transfected HEK293T cells were lysed in ice-cold RIPA buffer (50 mM Tris pH 7.5, 150 mM NaCl, 1% Triton, 0.1% SDS, 0.5% deoxycholic acid) with protease inhibitor cocktail (Roche). Samples were sonicated and then centrifuged at 14,000 rpm for 20 min at 4 °C. 5% of the supernatant was used as input and the remaining sample was incubated with Flag-beads or HA-beads for 2-4 hours for immunoprecipitation of 4xFlag-Rab33b, LULL1-HA, or GRASP55-HA.

All Western blot were performed according to standard procedure. Briefly, samples were boiled and separated by gel electrophoresis with 8 or 12% SDS-PAGE gels and transferred to a PVDF membrane. Membranes were incubated for 1 hour in 5% milk (Carnation) in TBST. Milk was washed and membranes were incubated overnight with indicated primary antibodies. Secondary antibodies were used were Alexa flour donkey anti-mouse 680 (Invitrogen) or donkey anti-rabbit 800 (Invitrogen) at 1:10,000 dilutions.

### Yeast culture and toxicity Assays

BY4741 yeast strain was used in all the experiments to test the toxicity of *Legionella* proteins. Plasmids expressing *Legionella* proteins were transformed into yeast cells with the standard lithium acetate procedure [10]. Cells were cultured to saturation in complete supplement mixture -Ura or -Ura-Leu media (Sunrise Science Products) containing 2% glucose. Cultured cells were then diluted to 0.3 OD600 and regrown to log phase before a 10-fold serial dilution with a starting density of 0.3 OD600. Dilutions were spotted on selective media containing either 2% glucose for repression, or 2% galactose for induction of pGal genes. Plates were incubated at 30°C for 3 days and then imaged using a BioRad ChemiDoc XRS imager. All yeast toxicity assays were performed in triplicate. The expression of *Legionella* proteins was confirmed by Western blotting of cell lysates prepared with glass bead agitation in lysis buffer (50 mM Tris pH 7.5, 150 mM NaCl, 0.2% Tergitol, 1 mM PMSF and Roche Protease Inhibitors).

### *Legionella* strains and infection

Strains of L. pneumophila used include the wild type Lp02 [11], the Dot/Icm deficient Lp03 [11]. The `Δ`*dupA* strain was created with triparental mating of the donor *E. coli* DH5αλpir carrying the suicide plasmid pSR47s-*dupA*, the pHelper strain, and the recipient Lp02 strain [9]. Integration of the plasmid was selected for with CYET plates containing 20 μg/mL of Kanamycin and counter selected on CYET containing 5% sucrose. Colonies were screened with PCR to identify genomic deletions. `Δ`*dupB* strain was obtained from Dr. Zhao-qing Luo (Purdue University). The `Δ`*dupA* `Δ`*dupB* strain was created using `Δ`*dupB* as a recipient strain. Complementation strains were produced by electroporation of pZL507 plasmids containing *dupA*, *dupB*, or their PDE mutants into the `Δ`*dupA* `Δ`*dupB* strain.

For *Legionella* infections, HEK293T cells were transfected with FCγRII and 4xFlag-Rab33 or LULL1-HA or GRASP55-HA for 24 hrs. Bacteria of indicated *Legionella* strains were opsonized with rabbit anti-legionella antibodies (1:500) at 37°C for 20 min before infection. The HEK293T cells were infected with post-exponential *L. pneumophila* strains at an MOI of 2 (for confocal imaging), or 10 (for Western blot) for the indicated amount of time.

### Intracellular growth assay

Intracellular growth of *L. pneumophila* was assayed using *A. castellanii* as a host cell. *A. castellanii* was propagated using PYG medium. Cells were grown to near confluency and then plated into 24-well culture dishes at a density of one million per well. The following day the cells were washed and equilibrated at 37 °C for 1 hour in *A. castellanii* buffer [12]. The amoebae were infected with stationary phase *L. pneumophila* cells at a multiplicity of infection (MOI) of 0.14 for 1 hour, washed one time to remove extracellular bacteria, and incubated for two days. *L. pneumophila* growth was assayed at 0, 18, 28, and 40 hours by lysing the infected amoebae with 0.05% saponin (Sigma, St. Louis, MO) in PBS and plating serially dilutions onto CYE plates. All growth assays were performed in triplicate.

### SILAC labeling and sample preparation

HEK293T cells were grown for 2 weeks in complete media containing normal lysine and arginine (“light”) or [13C6,15N2] lysine and [13C6,15N4] arginine (“heavy”, Sigma) prior to transfection. The labeled cells were passaged at 20% density into 6-well plates and were transfected with pCDNA3 HA-Ub G76A plasmid 24 hrs prior to infection. The cells grown in light medium were infected with the *`Δ`4sidE Legionella* strain and the cells grown in heavy medium were infected with the `Δ`*dupA* `Δ`*dupB* strain for 2 hrs. Cells were then washed two times with cold PBS and collected with lysis buffer (1% triton-X 100 in 50 mM Tris pH 8.0, 150 mM NaCl, 0.1% deoxycholate, phosphatase inhibitor cocktail, and protease inhibitor (Roche)). Cells were briefly sonicated (10% amplitude, 5 sec duration, pulse) and centrifuged at 14,000 rpm for 15 minutes at 4°C to remove insoluble fraction. Supernatant was collected and mixed with 30 µL slurry of EZ-view Red anti-HA resin. The mixture was incubated on a nutating mixer at 4°C for 4 hrs. The anti-HA resins were briefly centrifuged down and subsequently washed with washing buffer (1% triton-X in 50 mM Tris pH 8.0 and 150 mM NaCl) for a total of 5 washes. Bound proteins were eluted by incubating of the resin with 45 µL of elution buffer containing 1% SDS in 100 mM Tris pH 8.0 at 65°C for 15 minutes. Eluted proteins from light or heavy media grown cells were mixed together, reduced, alkylated with iodoacetamide and then precipitated with three volumes of a solution containing 50% acetone and 50% ethanol. Precipitated proteins were solubilized in a solution of 2 M urea, 50 mM Tris-HCl pH 8.0, and 150 mM NaCl, and then digested with Pierce trypsin protease MS-grade (Thermo Scientific) overnight at 37°C. Digested peptides were acidified with 0.2% Trifluoroacetic acid and formic acid and then desalted with Sep-Pak C18 column (Waters) for mass spectrometry analysis.

### Mass spectrometry data acquisition and analysis

Samples resuspended in 0.1% trifluoroacetic acid were subjected to LC-MS/MS analysis using a 20-cm-long 125-µm inner diameter column packed in-house with 3 µm C18 reversed-phase particles (Magic C18 AQ beads, Bruker). Separated peptides were electrosprayed into a Q-Exactive HF Orbitrap mass spectrometer (Thermo Fisher Scientific). Xcalibur software (Thermo Fischer Scientific) was used for the data acquisition and the Q-Exactive was operated in data-dependent mode. Survey scans were acquired in the Orbitrap mass analyzer over the range of 380 to 2000 m/z with a mass resolution of 70.000 (at m/z 200). MS/MS was performed selecting up to the 10 most abundant ions with a charge state of 2, 3 or 4 within an isolation window of 2.0 m/z. Selected ions were fragmented by Higher-energy Collisional Dissociation (HCD) with normalized collision energy of 27 and the tandem mass spectra was acquired in the Orbitrap mass analyzer with a mass resolution of 17.500 (at m/z 200). Repeat selection of peptides was kept to a minimum by dynamic exclusion of the sequenced peptides for 30 seconds. PSM searches were performed using the Sequest-based engine SORCERER (Sage N Research, Inc.). Digestion parameters were set to semi-tryptic. Differential modifications for the PSM search included of 8.0142 Daltons for lysine, 10.00827 Daltons for arginine, and a static mass modification of 57.021465 Daltons for alkylated cysteine residues. For a PSM to be considered for SILAC quantification, we required a mass accuracy of at least 15 ppms for the experimental and theoretical precursor ions and a peptide prophet score of >0.7. For a protein to be considered in the quantification, we required at least 6 independent PSMs and a standard deviation <5.5. For the identification of phosphoribosylated peptides and localization of sites was obtained from MaxQuant software (v.1.6.3.4) using the *Legionella pneumophila* proteome (2932 entries, Uniprot) incorporated with the ubiquitin-40S ribosomal protein S27a sequence. Search criteria consisted in fixed modifications for carbamidomethyl cysteines (57.02146 Da), variable modification for phosphorybosylated arginines (212.00859 Da) and score cut-of at a maximum of 1% false discovery rate based on decoy (reversed) database searches.

---

**Supplemental Fig. S1.**
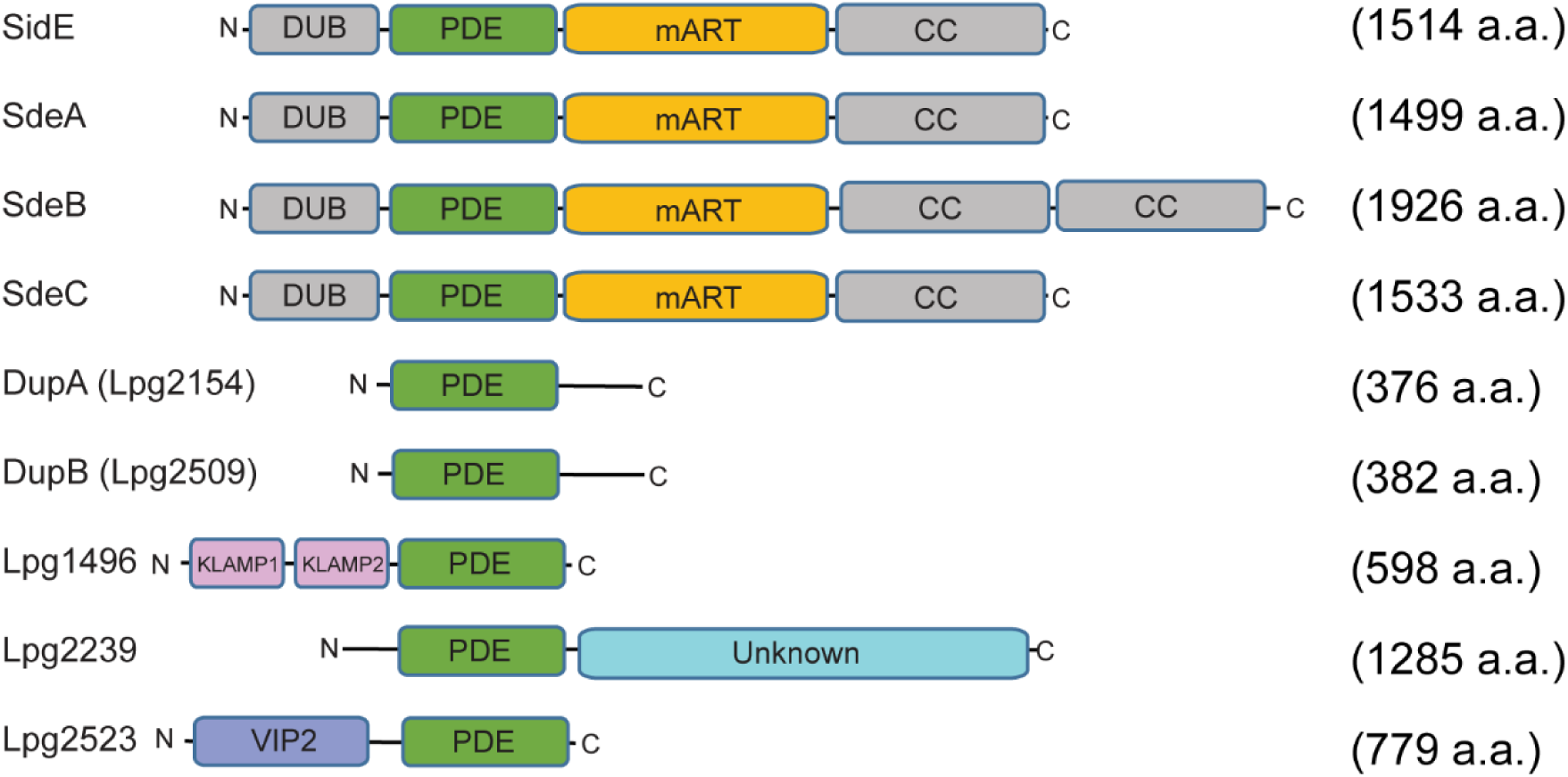
Schematic diagram of the PR-ubiquitination reaction and PDE domain-containing *L. pneumophila* effectors. (*B*) Domain organization of PDE domain-containing *L. pneumophila* effectors. It is notable that the PDE is highly conserved between DupA and DupB [1], while there is no detectable homology between their ∼40 amino acids long C-terminal tails. Functional or structural domains are labeled. DUB: deubiquitinase domain; PDE: phosphodiesterase; mART: mono-ADP ribosyltransferase, CC: coiled-coil; KLAMP: kinase-like ATP binding region-containing protein and S-adenosylmethionine-dependent methyltransferase protein; VIP2: vegetative insecticidal protein 2.

**Supplemental Fig. S2.**
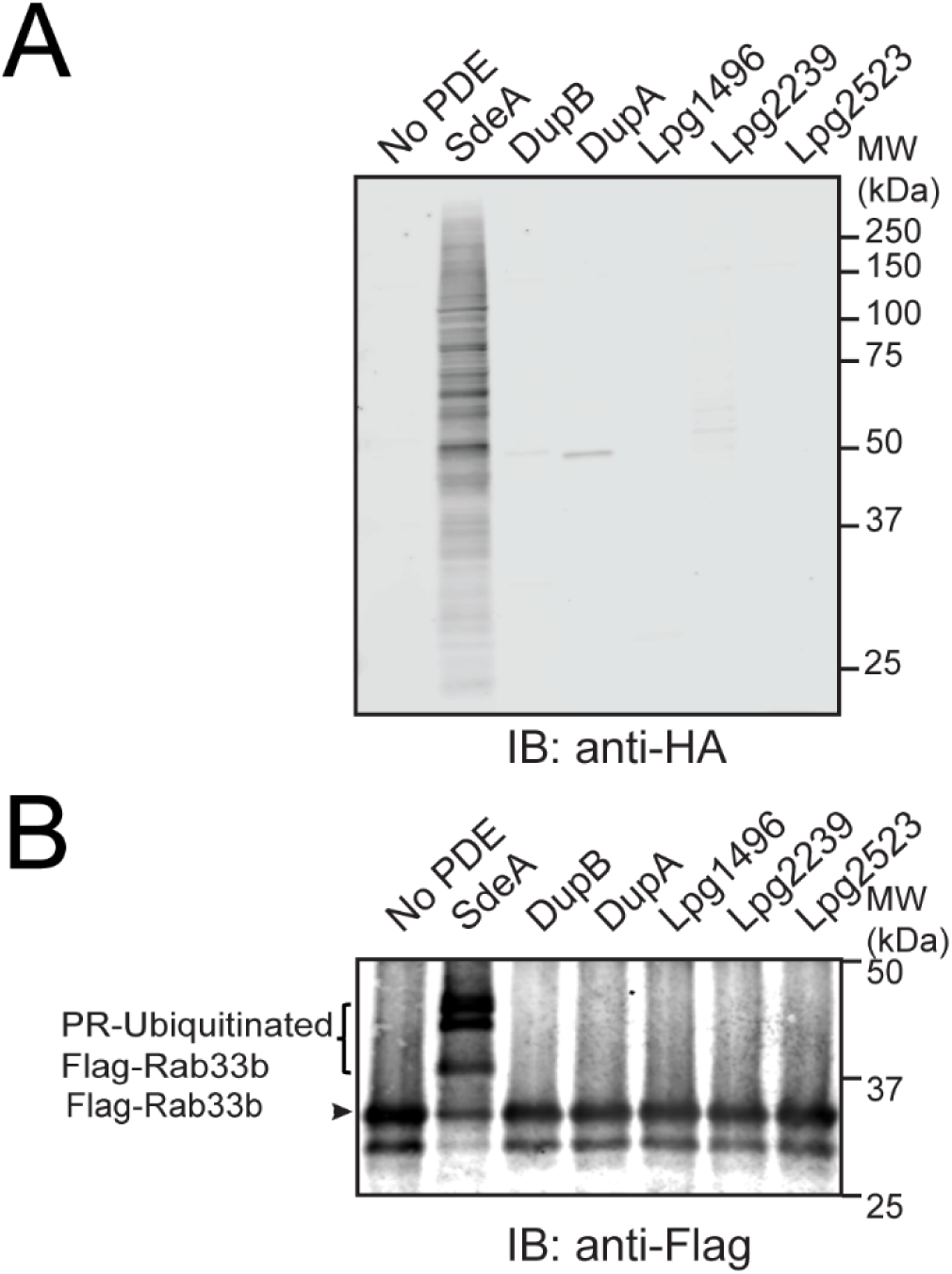
PR-Ubiquitination ligation assay of the PDE domains from the specified *Legionella* effectors. (*A*) HEK293T whole cell lysates were mixed with HA tagged ADPR-Ub and then treated with recombinant PDE domain proteins from indicated *Legionella* effectors. PR-ubiquitinated species were detected by immunoblot against HA. (*B*) Recombinant 4xFlag-Rab33b proteins were incubated with ADPR-Ub and PDE domain proteins from indicated *Legionella* effectors. PR-ubiquitination of Rab33b was monitored by immunoblot against Flag.

**Supplemental Fig. S3.**
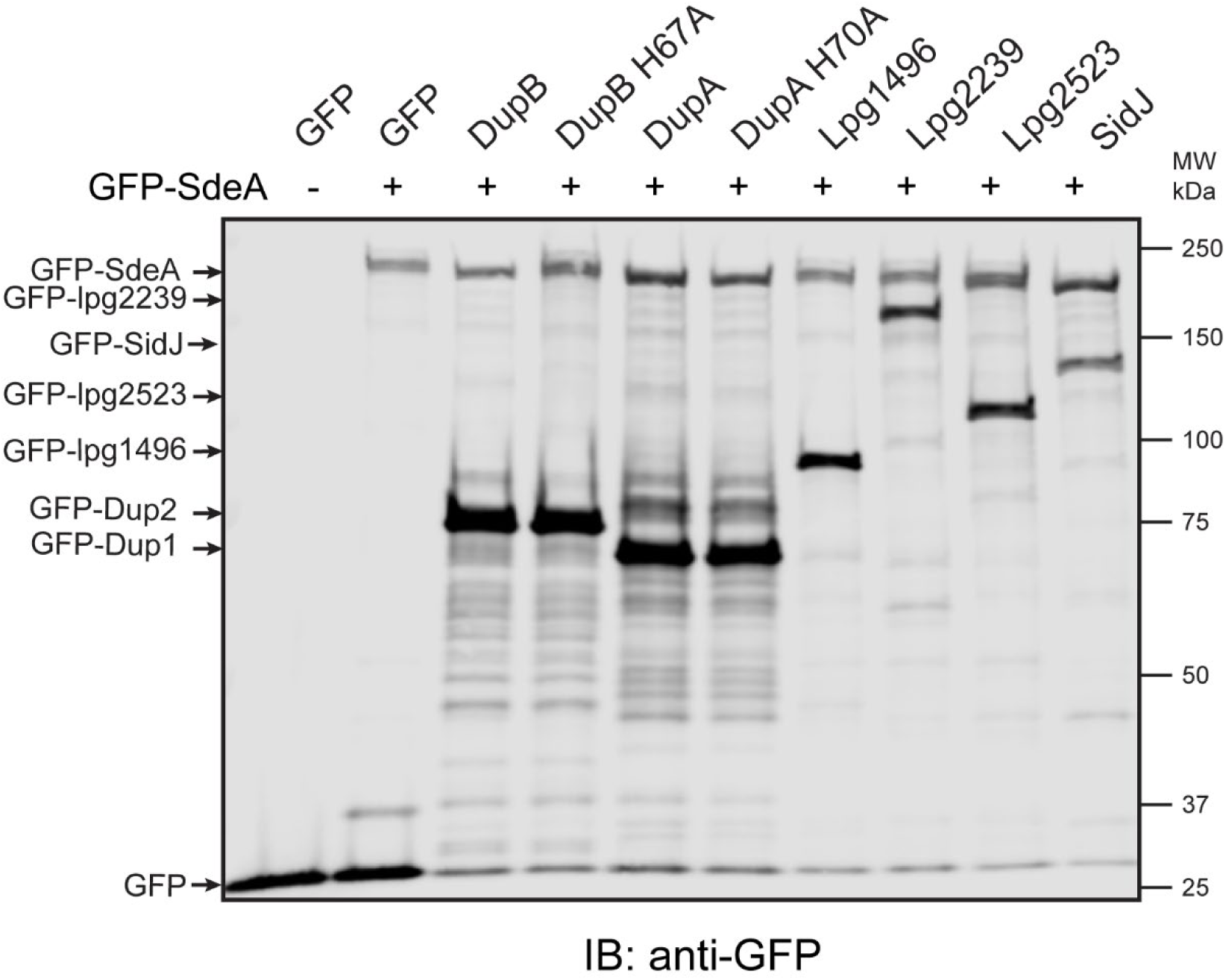
4xFlag-Rab33b and EGFP-SdeA were co-expressed with full-length protein of indicated EGFP-tagged *Legionella* effector in HEK293T cells. GFP or GFP-tagged proteins were analyzed by Western blot against anti-GFP.

**Supplemental Fig. S4.**
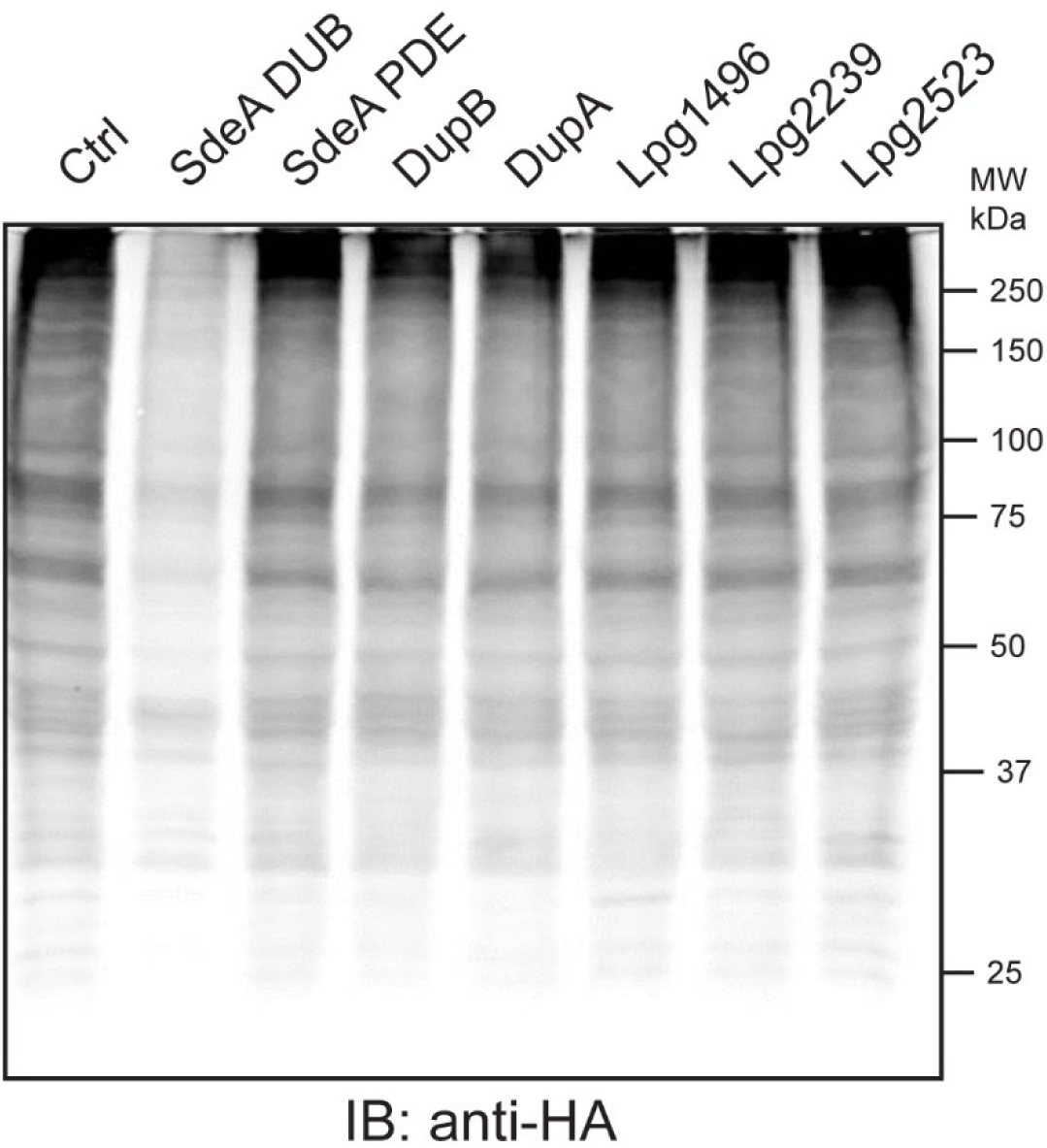
DupA and DupB do not cleave canonical poly-ubiquitinated conjugates. Cell lysates from HEK293T cells expressing HA-Ub were treated with either 0.2 μM SdeA DUB domain or 1 μM protein of indicated PDE domains. The reaction products were analyzed by Western blot against HA.

**Supplemental Fig. S5.**
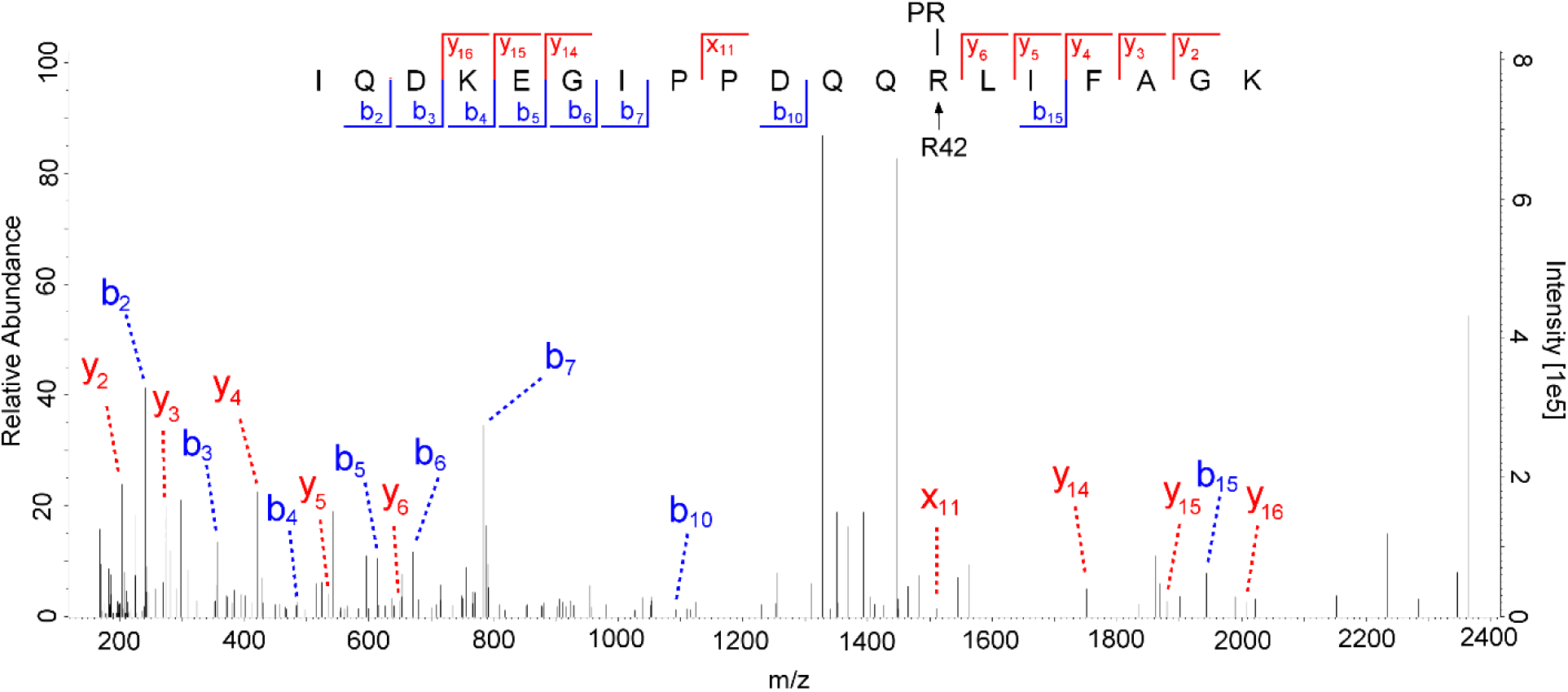
MS/MS spectrum produced by HCD fragmentation of a Ub peptide containing the phosphoribosyl modification at Ub R42. Recombinant 4xFlag-Rab33b was PR-ubiquitinated by SdeA-Core and purified with anti-Flag beads. The bound proteins were then treated with recombinant DupB PDE domain, and the supernatant processed and analyzed by LC-MS/MS in triplicate. The spectrum represents a peptide (MH+ = 2365.194) with the indicated amino acid sequence mass plus the addition of 212.009 Da (phosphoribosilation) at a high confidence mass accuracy of 1ppm. When submitted to Higher-energy collisional dissociation (HCD), the fragmented ions labeled in red (c-terminal fragments) and blue (n-terminal fragments) supports the presence of the mass shift of 201.009 Da between Q40 and R42, which corresponds to the phosphoribosyl modification at Ub R42.

**Supplemental Fig. S6.**
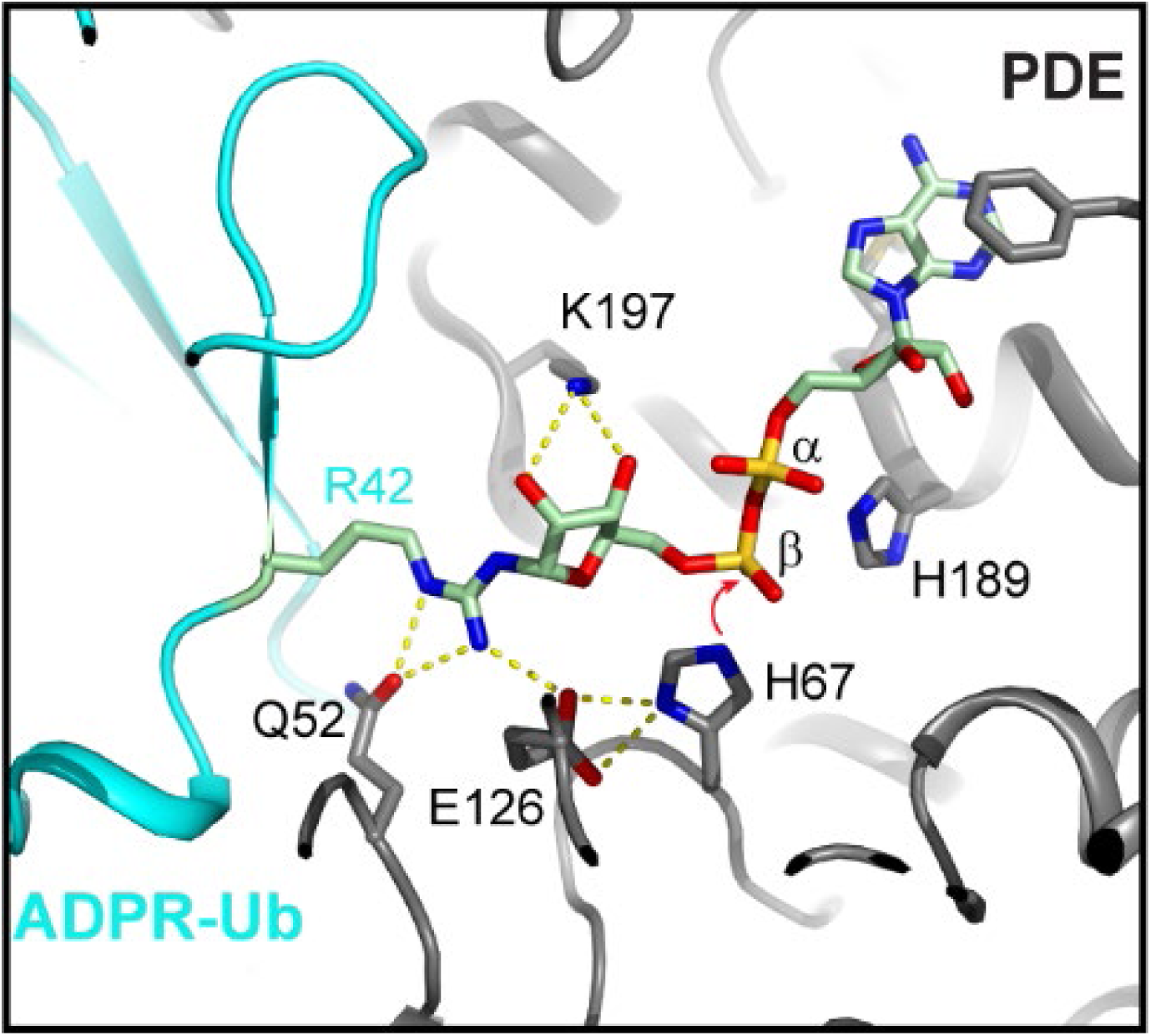
Ribbon diagram of complex structure of ADPR-Ub with the PDE domain of DupB (PDB ID: 6B7O). The β phosphate of the ADPR moiety is positioned in close proximity to the catalytic histidine H67. The complex structure indicates that during the cleavage of ADPR-Ub, H67 attacks the β phosphate to break the phosphodiester bond and generate PR-Ub and H189 provides a proton to release AMP.

**Supplemental Fig. S7.**
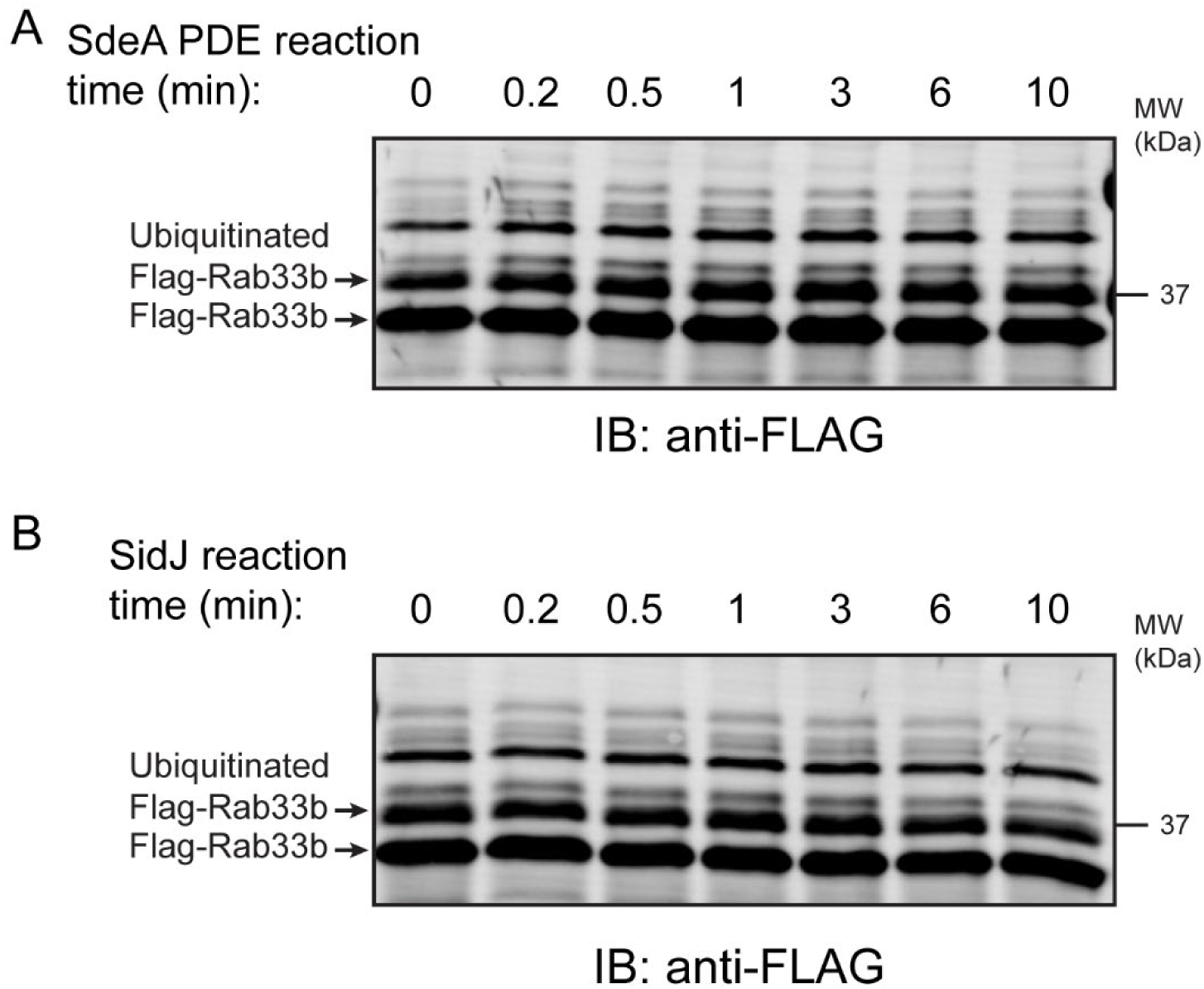
The reaction time course of the cleavage of PR-ubiquitinated Rab33b by the PDE domain of SdeA and SidJ. (*A*) PR-ubiquitinated Rab33b was generated by incubating 4 μM 4xFlag-Rab33b with 1 μM SdeA-Core, 25 μM HA-Ub and 1 mM NAD^+^ for 1 hr at 37 °C. Then the product was mixed with cobalt resin to remove 6xHis-sumo-SdeA-Core and the supernatant was treated with 1 μM SdeA PDE domain for the indicated periods of time at 37 °C. The final reaction products were analyzed by Western blot using anti-Flag antibodies. (*B*) Similar reactions with SidJ (1 μM SidJ 89-853, 5 mM MgCl2, 5 mM Calmodulin, 5 mM Glutamate, 1 mM ATP).

**Supplemental Fig. S8.**
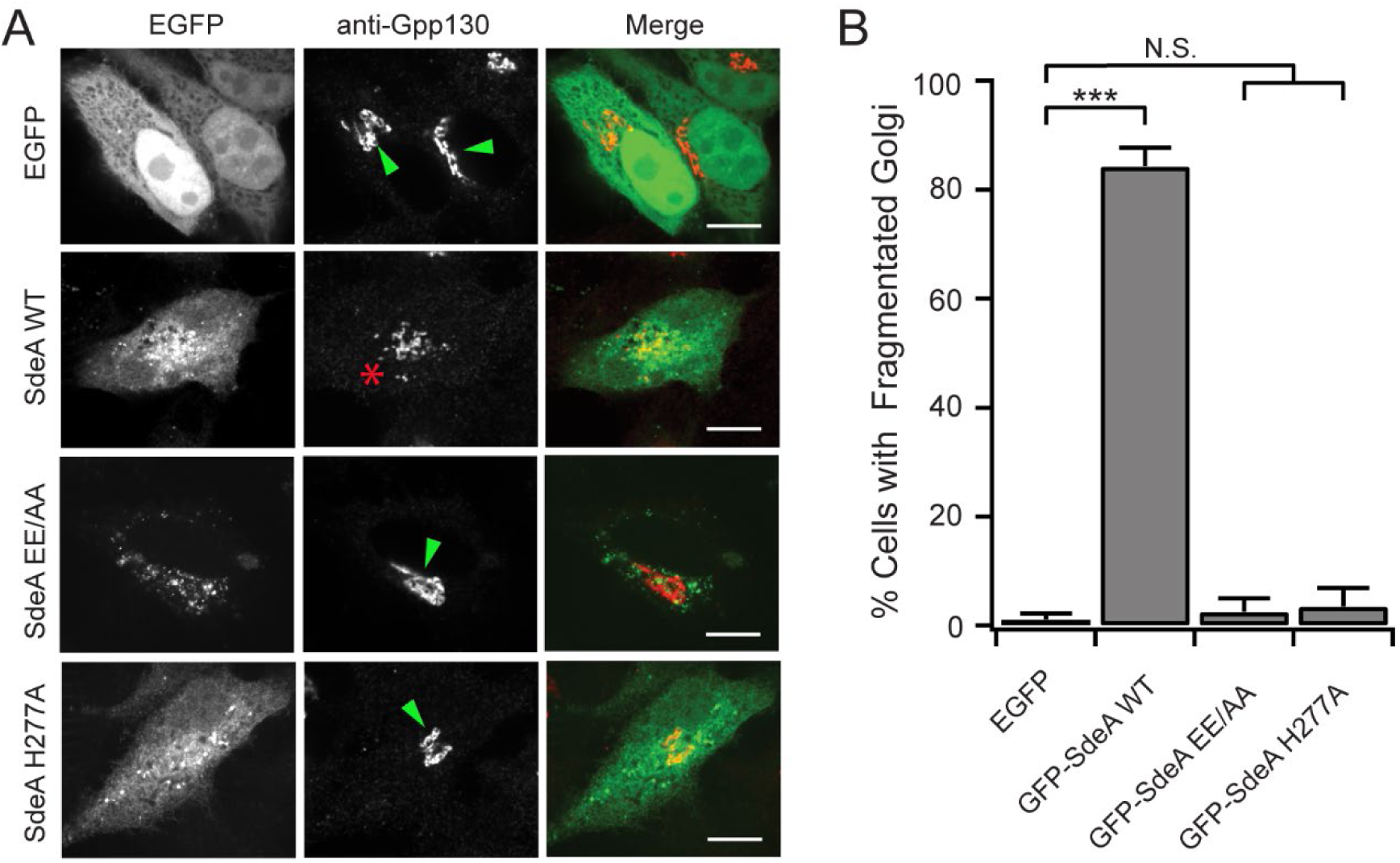
The PR-ubiquitination activity of SdeA induces dispersed Golgi phenotype. (*A*) Representative images showing the Golgi morphology of HeLa cells expressing EGFP or SdeA wild-type or mutants for 20 hours. Cells were fixed and immunostained with rabbit anti-Gpp130. Transfected cells with normal and dispersed Golgi structures were marked with green arrow heads or red stars, respectively. Scale Bar: 10 μm. (*B*) Quantification of the dispersed Golgi phenotype for the cells in (A). Data are shown as means ± SEM of three independent experiments. More than 40 cells for each condition were counted. ***P<0.001, N.S.: Not Significant.

**Supplemental Fig. S9.**
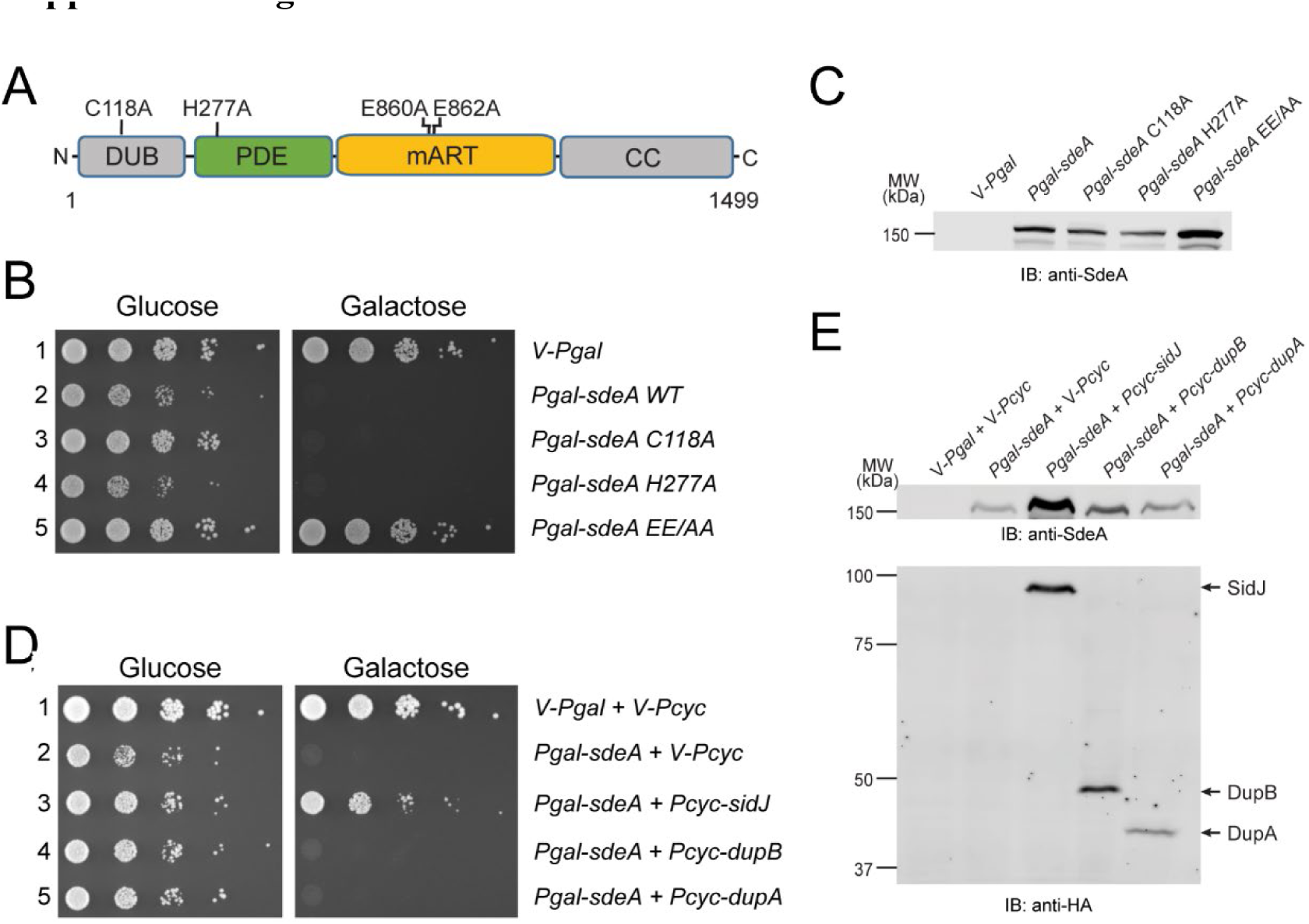
DupA and DupB fail to suppress SdeA yeast toxicity under the control of a low expression constitutive promotor. (*A*) Domain architecture of SdeA displaying catalytic mutants used in SdeA yeast toxicity assay. (*B*) Yeast strains expressing vector control or SdeA catalytic point mutants were serially diluted and spotted onto glucose and galactose medium. Images were acquired 3 days after incubation at 30 °C. (*C*) Cell lysates were prepared from yeast cells expressing SdeA or SdeA catalytic mutants after galactose induction for 8 hrs. The expression of SdeA proteins was verified by Western blot using anti-SdeA antibody. (*D*) Serially diluted yeast strains expressing vector control or SdeA and coexpressed with SidJ, DupA, or DupB were spotted and grown on glucose and galactose medium. The expressing of SidJ, DupB, and DupA was under the control of the low constitutive *Pcyc* promotor. (*E*) Yeast cell lysates expressing galactose induced SdeA and low constitutive *Pcyc* expressed HA tagged SidJ, DupB or DupA were verified by Western blot using anti SdeA (top panel) or anti-HA (bottom panel).

**Supplemental Fig. S10.**
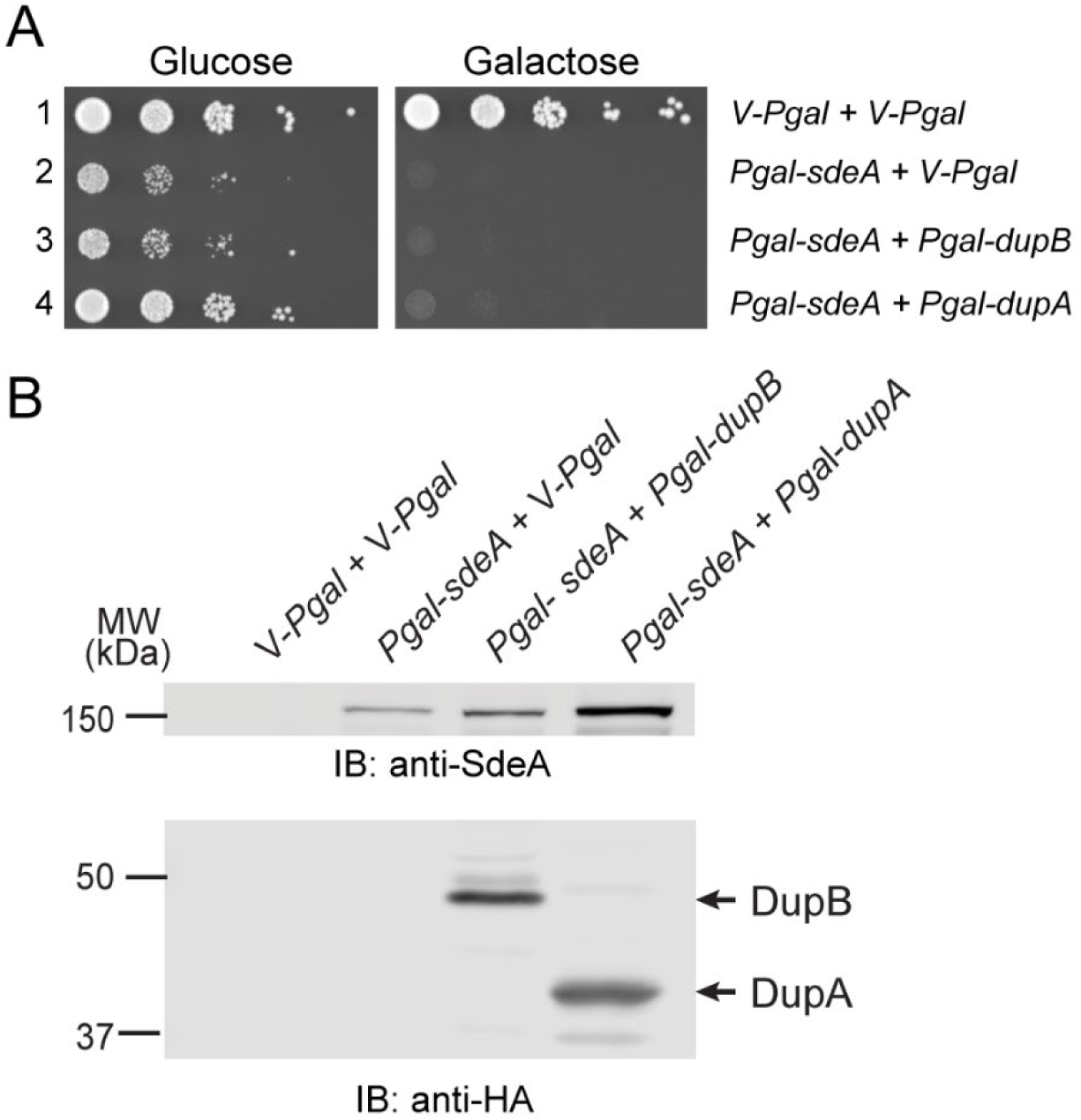
DupA and DupB fail to suppress SdeA yeast toxicity under the control of a high expression inducible promotor. (*A*) Serially diluted yeast strains expressing vector control, SdeA, DupA, or DupB were spotted on glucose and galactose medium. The expression of DupA, DupB, and SdeA was under the control of inducible *Pgal* promotor. (*B*) Yeast cell lysates expressing galactose induced SdeA and HA tagged DupB or DupA were verified by Western blot using anti SdeA (top panel) or anti-HA (bottom panel).

**Supplemental Fig. S11.**
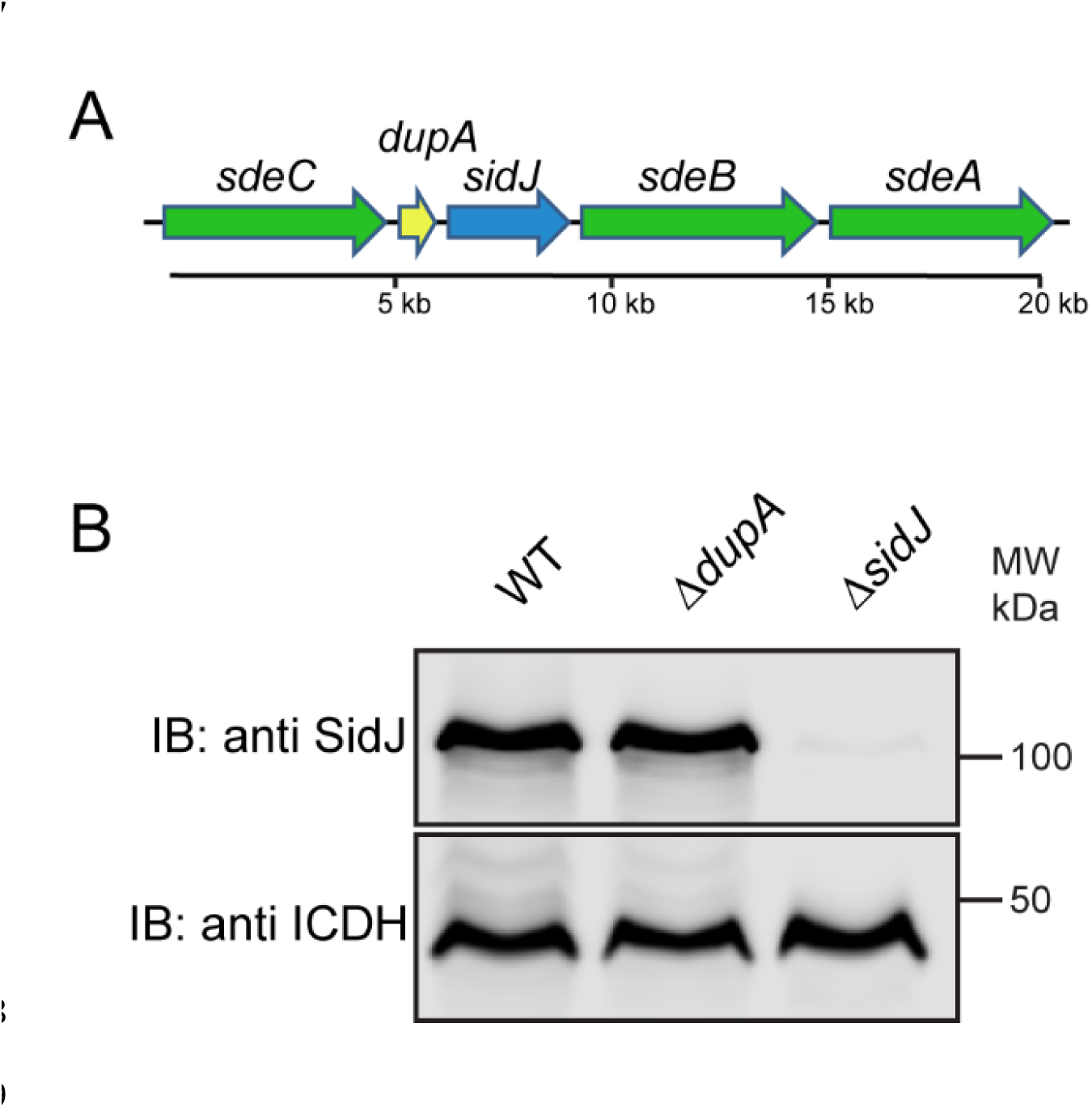
*dupA* deletion has no polar effect on the expression of its adjacent gene *sidJ*. (*A*) The genome locus of *dupA*. This locus is comprised of genes encoding three PR-Ub ligases (SdeA, SdeB, and SdeC), the PR-Ub deubiquitinase DupA, and SidJ. (*B*) Western blot analysis of the expression of SidJ. Bacterium of indicated *Legionella* strains was grown overnight in AYET medium and lysed with 4% SDS. SidJ was visualized by Western blot using anti-SidJ antibodies (top panel). Western blot of isocitrate dehydrogenase (ICDH) with anti-ICDH antibodies to serve as protein loading control (bottom panel).

**Supplemental Fig. S12.**
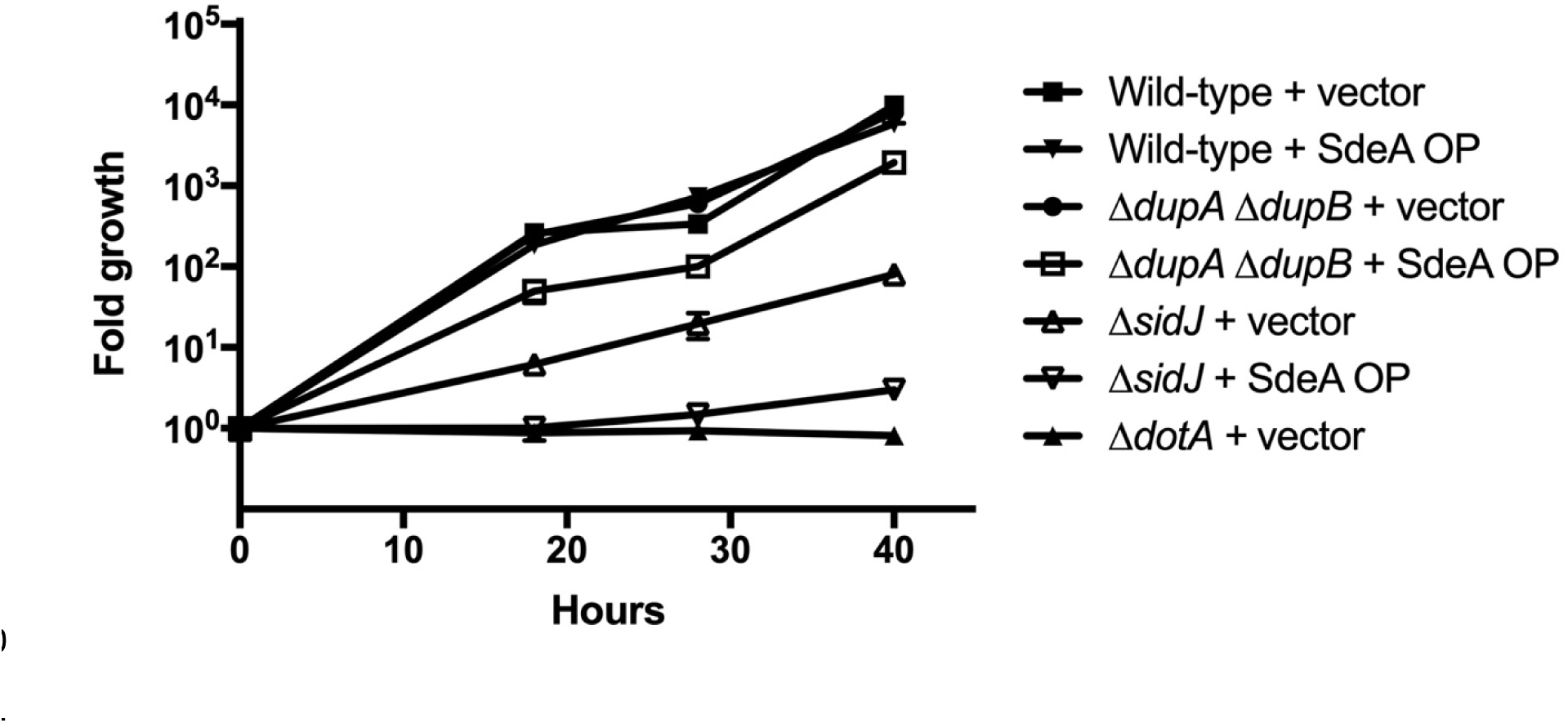
In contrast to the Δ*sidJ* mutant, the Δ*dupA* Δ*dupB* mutant does not have a growth defect within *Acanthamoebae castellanii* but both mutants show diminished growth by over-expression of SdeA. A wild-type *Legionella* strain (JV1139), a wild-type strain over-expressing SdeA (JV6445), a Δ*dupA* Δ*dupB* mutant (JV9336), a Δ*dupA* Δ*dupB* over-expressing SdeA (JV9336), a Δ*sidJ* mutant (JV4925), and a Δ*sidJ* mutant over-expressing SdeA (JV6451) were used to infect *A. castellanii* cells. Growth was assayed by plating for colony forming units (CFUs) over 40 hours. Data is representative of three independent experiments. There is a statistical difference (P < 0.05) between the growth of the wild-type strain and the Δ*dupA* Δ*dupB* over-expressing SdeA strain as compared by Student’s t test at 18 hours (P = 0.03), 28 hours (P < 0.0001) and 40 hours (P < 0.001).

**Supplemental Fig. S13.**
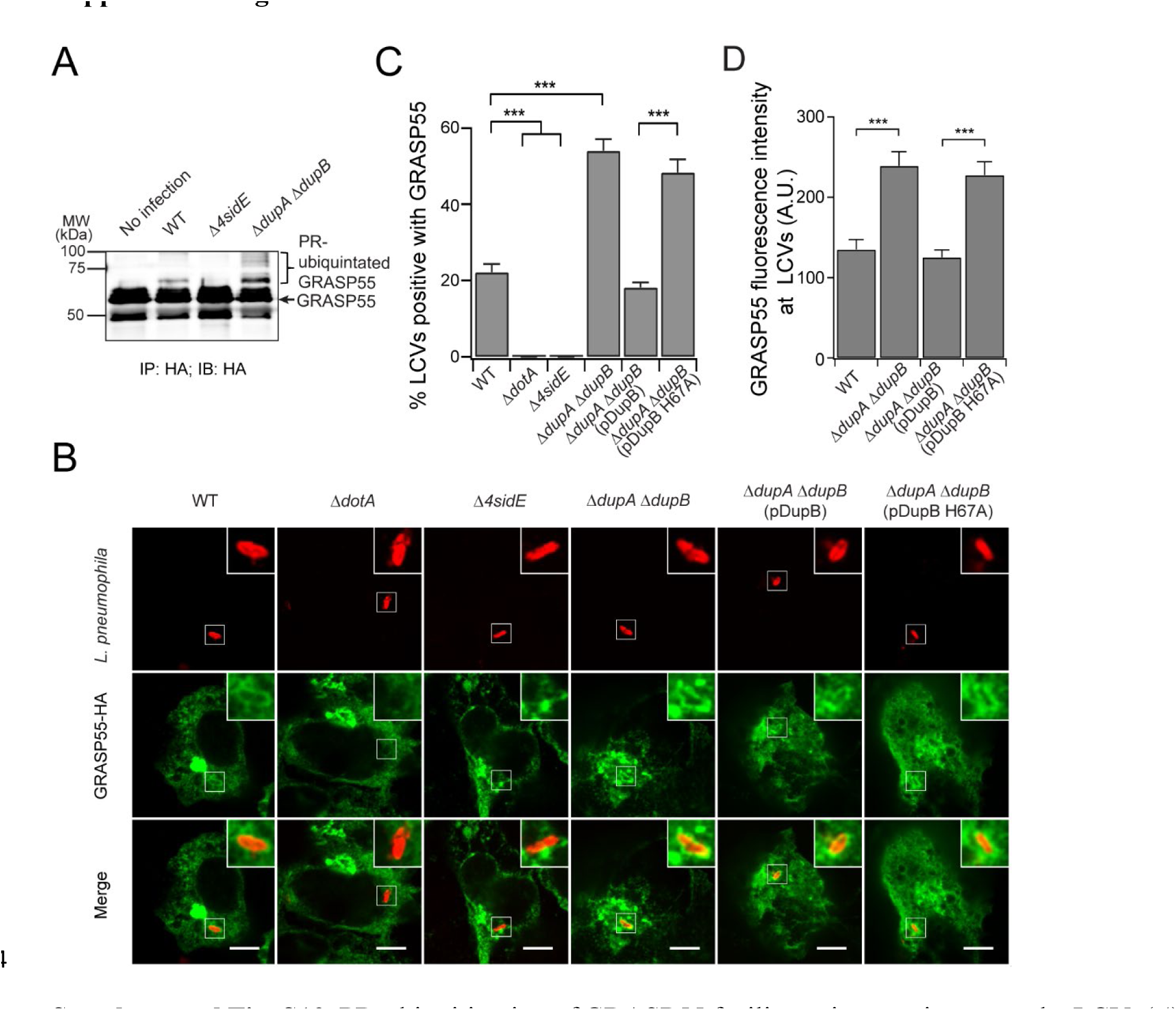
PR-ubiquitination of GRASP55 facilitates its recruitment to the LCV. (*A*) GRASP55 was PR-ubiquitinated upon *Legionella* infection. HEK293T cells were transfected with GRASP55-HA and then infected with indicated *Legionella* strains for 2 hrs. GRASP55 was enriched by anti-HA immunoprecipitation and analyzed by anti-HA Western blot. (*B*) Representative confocal images show the recruitment of GRASP55 (green) in HEK293T cells challenged with specified *Legionella* strains (red) for 2 hrs. Scale Bar: 5 μm. (*C*) Quantitative analysis of GRASP55-positive LCVs in cells infected with the indicated *Legionella* strains. Data were shown as means ±SEM of three independent experiments. More than 80 LCVs were counted for each condition. ***P<0.001. (*D*) Quantitative analysis of the GRASP55 fluorescence intensity associated with the LCV. Data were shown as means ±SEM of three independent experiments. The value was averaged from more than 18 GRASP55-positve LCVs for each condition. ***P<0.001.

